# An end-to-end deep learning method for rotamer-free protein side-chain packing

**DOI:** 10.1101/2022.03.11.483812

**Authors:** Matt McPartlon, Jinbo Xu

## Abstract

Protein side-chain packing (PSCP), the task of determining amino acid side-chain conformations, has important applications to protein structure prediction, refinement, and design. Many methods have been proposed to resolve this problem, but their accuracy is still unsatisfactory. To address this, we present AttnPacker, an end-to-end, SE(3)-equivariant deep graph transformer architecture for the direct prediction of side-chain coordinates. Unlike existing methods, AttnPacker directly incorporates backbone geometry to simultaneously compute all amino acid side-chain atom coordinates without delegating to a rotamer library, or performing expensive conformational search or sampling steps. Tested on the CASP13 and CASP14 native and non-native protein backbones, AttnPacker predicts side-chain conformations with RMSD significantly lower than the best side-chain packing methods (SCWRL4, FASPR, Rosetta Packer, and DLPacker), and achieves even greater improvements on surface residues. In addition to RMSD, our method also achieves top performance in side-chain dihedral prediction across both data sets.

## 1 Introduction

Protein side-chain packing (PSCP) involves predicting the three dimensional coordinates of side-chain atoms given backbone conformation (coordinates) and primary sequence. This problem has important applications to protein structure prediction [11, 10, 6], design [23, 26, 8, 29], and protein-protein interactions [28, 14]. Traditional methods for PSCP rely on minimizing some energy function over a set of rotamers from a fixed library [15, 34, 25, 4, 1, 9, 32, 5]. These methods tend to differ primarily in their choice of rotamer library, energy function, and energy minimization procedure. Although many of these methods have shown success, the use of search heuristics coupled with a discrete sampling procedure could ultimately limit their accuracy. Currently, the fastest methods (OSCAR-star[25], FASPR[15], SCWRL4[9]) do not employ deep learning (DL), and are rotamer library-based.

Several machine learning (ML) methods have been proposed for the task of side-chain prediction [22, 21, 30, 31, 34, 20, 32]. One of the earliest of these methods, SIDEPro [22], attempts to learn an additive energy function over pairwise atomic distances for each side-chain rotamer. This is achieved by training a family of 156 feedforward networks - one for each amino acid and contacting atom type. The rotamer with the lowest energy is then selected. DLPacker[21] formulates packing as an image-to-image transformation problem and employs a deep U-net-style neural network. The method iteratively predicts side-chain atom positions using a voxelized representation of the residue’s local environment as input, and outputs densities for the respective side-chain’s atoms. The output densities are then compared to a rotamer database and the closest matching rotamer is selected. The most recent version of OPUS-Rota4[31] uses a pipeline of multiple deep networks to predict side-chain coordinates. The method uses predicted side-chain dihedral angles to obtain an initial model, and then applies gradient descent on predicted distance constraints to obtain a final structure. It is worth noting that OPUS-Rota4, to the best of our knowledge, is the only ML-based PSCP method that directly utilizes MSA (multiple sequence alignment) as part of its input.

Here we present AttnPacker, a new deep architecture for PSCP. Our method is inspired by recent breakthroughs in modelling three-dimensional data, and architectures for protein structure prediction - most notably AlphaFold2[18], Tensor Field Networks (TFN) [27], and SE(3)-Transformer[12]. By modifying and combining components of these architectures, we are able to significantly outperform traditional PSCP methods as well as machine learning methods in terms of speed, memory efficiency, and overall accuracy using only features derived directly from primary sequence and backbone coordinates as input.

Specifically, we introduce deep graph transformer architectures for updating both residue and pairwise features based on the input protein’s backbone conformation. Inspired by AlphaFold2, we propose *locality-aware triangle updates* to refine our pairwise features using a graph-based framework for computing triangle attention and multiplication updates. By doing this, we are able to significantly reduce the memory required for performing triangle updates which enables us to build higher capacity models. In addition, we explore several SE(3)-equivariant attention mechanisms and propose an SE(3)-equivariant transformer architecture for learning from 3D points.

Our method, AttnPacker, significantly outperforms traditional PSCP methods on CASP13 and CASP14 native backbones with average reconstructed RMSD over 19% and 25% lower than the next best method on each test set. AttnPacker also surpasses deep learning DLPacker, with 13% lower average RMSD on the each test set. In addition, our method achieves top RMSD scores for CASP14 non-native backbone targets predicted by AlphaFold2. On top of our method’s favorable performance in RMSD minimization, it also achieves the lowest χ_1–4_ mean absolute error (MAE) across all methods on the CASP13 and CASP14 targets.

## 2 Methods

### 2.1 Network Architecture

Our method draws on several recent breakthroughs in deep learning and protein structure prediction, namely the two-stage architecture introduced by AlphaFold2, and a TFN-based equivariant transformer inspired by Fuchs et al[12]. Moreover, our model directly predicts the 3D coordinates of all side-chain atoms for a given protein using only backbone coordinates and primary sequence as input. A detailed overview of input features and the input embedding procedure can be found in Supplementary Material, Table S3, and Figure S1.

The first component, outlined in Figure 1, is a deep graph transformer network which utilizes the geometry of the input backbone to revise node and pair features. This component is similar to the AlphaFold2’s ’Evoformer’ module for processing MSA and pair features, but we replace axial self-attention and global triangle updates with graph based self-attention and triangle updates for our residue and pair features. In AlphaFold2, triangle updates are performed by considering all triples of residues in the input protein, updating the corresponding pair representations in a geometrically motivated manner. These procedures require *O* (*L*^3^*d*) time to compute, and Ω (*L*^3^) space to store attention logits where *L* is the number of residues and *d* is the hidden dimension of pair features. In our setting, we are able to avoid this restrictive time and space complexity by incorporating locality information derived from the protein’s backbone atom coordinates. To do this, we introduce locality aware triangle updates (Section 2.2) and compare the performance of this approach against the update procedures used in AlphaFold2.

**Figure 1:**
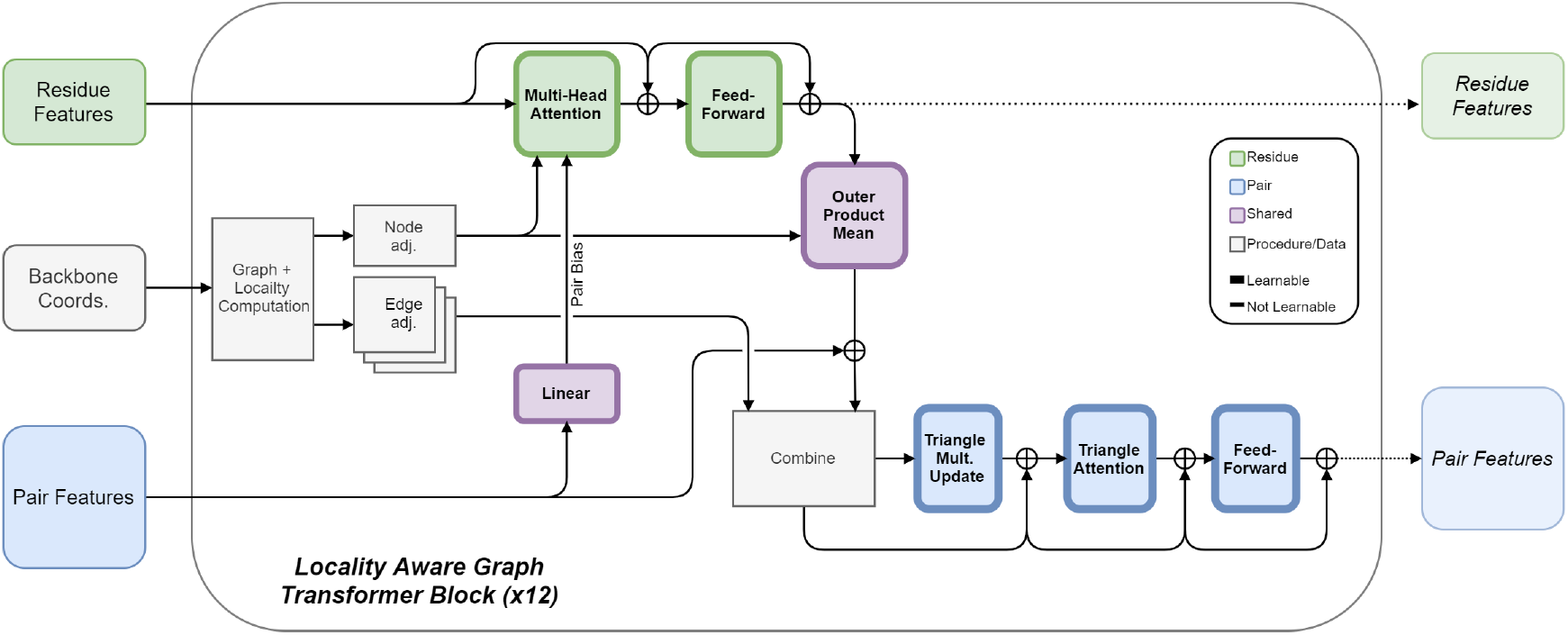
Architecture of Locality Aware Graph Transformer Block. Standard multi-headed graph attention is used to update node features, while edges between adjacent nodes are updated with triangle multiplication and attention. For each triangle update, edge adjacency information is used to select only those triangles having maximum side-length less than a fixed threshold.

The second component of our architecture is an SE(3)-equivariant neural network which takes the output of the first component, along with the protein’s backbone *C_α_*-coordinates, to simultaneously predict each residue’s sidechain coordinates. Our equivariant architecture is derived from the SE(3)-Transformer introduced by Fuchs et al. Similar to the SE(3)-Transformer, our attention blocks use TFNs to produce keys and values for scalar and point features. But our implementation differs in a few key areas. First, our implementation uses shared attention with dot-product based similarity (see Table 5). That is, we combine the attention logits of each feature type to produce shared attention weights. Furthermore, in each attention block, we augment the input to the TFN radial kernel with pairwise distance information, and make further use of the pair features to bias attention weights and update scalar features (see Algorithm S2, Algorithm S3). The architecture is outlined in Figure 2, full implementation details can be found in Section S6, and ablation studies guiding these decisions are given in, Table 6, and Table 7.

**Figure 2:**
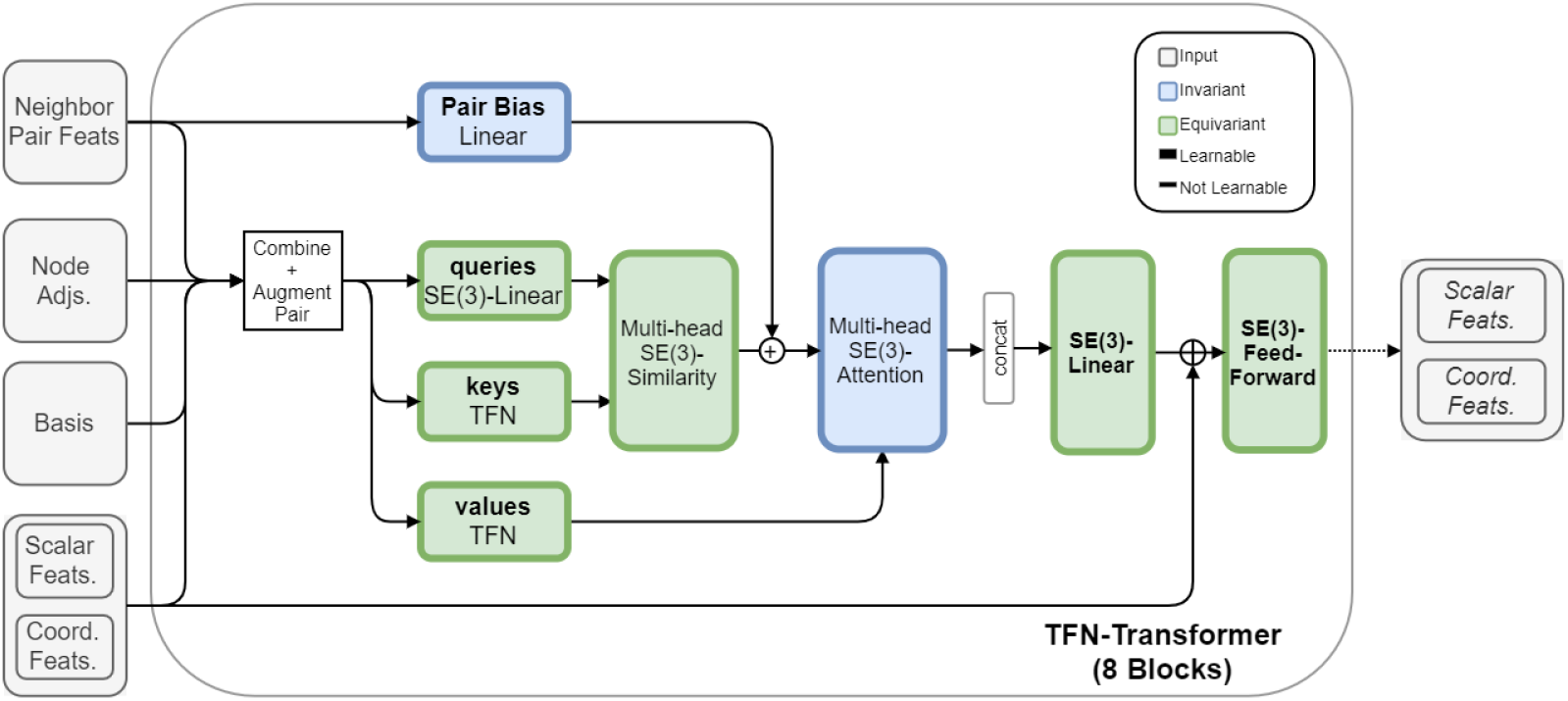
Architecture of our TFN-based transformer block. Keys and values are computed using TFNs. The input to each TFN radial kernel are pair features, along with distance information between points for the corresponding pair. Pair features are also linearly projected to bias the similarity logits for adjacent pairs of residues.

### 2.2 Locality Aware Triangle Updates

We experimented with multiple approaches for improving the time and space requirements of triangle updates. We ultimately settled on a hypergraph-based approach where a separate spatial graph is constructed for edge features. In this graph, there is a vertex *v_ij_* for residue pair *i, j* if the distance *d*(*i, j*) between the corresponding *C_α_* atoms is within a threshold *θ*. We add hyperedges between *v_ij_* and *v_ik_*, *v_jk_* only if max {*d*(*i, k*), *d*(*j, k*)} < *θ*. Interpreting this graph in the context of triangles, the vertices coincide with triangle edges of side length at most *θ*, and edges correspond to triangles. The process is illustrated in Figure 3.

**Figure 3:**
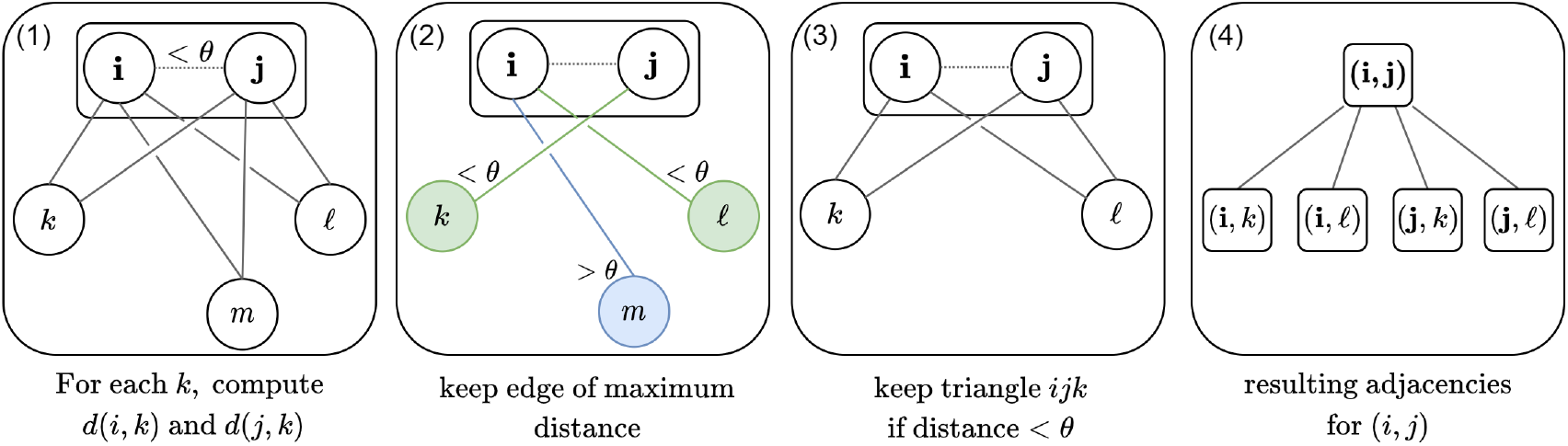
Overview of pair graph generation for locality aware attention. Each pair of residues within a fixed distance is included as a node in this graph. (A) We begin with the spatial graph for residues and focus on a specific pair (*i, j*). (B) For each node *x*, we then compute the maximum distance between (*i, x*) and (*j, x*) and disregard residues falling outside our distance threshold *θ*. (C) All triangles that include pair (*i, j*) in the remaining graph will have maximum side-length at most *θ*. (D) The final graph has nodes for each edge in (*C*), and edges for each triangle.

In order to efficiently compute triangle updates, we select the 2*N_e_* nearest neighbors of each vertex *v_ij_* using max {*d*(*i, k*), *d*(*j, k*)} as our distance function. This results in a 3-uniform, *N_e_*-regular hypergraph containing as many nodes as there are underlying residues with *C_α_* distance less than the threshold *θ*. We denote the latter quantity by *N_v_*. Triangle multiplication and attention updates are straight forward to apply since the neighbors of each node *v_ij_* span strictly over residues *k* for which the corresponding triangle is well-defined. The update operations require time and space proportional to *O*(*N_v_* · *N_e_*) in total. If the distance threshold *θ* is chosen so that the number of resulting pairs is linear in the number of atoms, then choosing constant *N_e_* results in *N_v_* · *N_e_* = *o*(*n*^2^) space complexity, and time complexity *O*(*N_v_* · *N_e_* · *d*). Computing the hypergraph still requires *O*(*L*^3^) time, but this can be performed on a CPU as a pre-processing step.

## 3 Results

We compare side-chain reconstruction accuracy for three of our models against several popular methods: DLPacker, RosettaPacker, SCWRL4, and FASPR. To understand the impact of different architectural components, we list results of our TFN-Transformer without triangle updates (*TFN*), with standard triangle updates (*+ Tri*), and with locality aware triangle updates (*+Tri+Local*), respectively. We use the same hyperparameters for the TFN-Transformer component, and use a distance cutoff of 15Å with 30 nearest neighbors for local triangle updates. All other hyperparameters are held constant unless otherwise specified. A full summary of model hyperparameters, training data, and training procedure can be found in Table S1, and Section S3.

To evaluate the performance of each method, we consider residue-level RMSD and dihedral angle deviations between the predicted and native side-chains. RMSD between predicted and native side-chain atoms is computed separately for each residue, and averaged over all residues in the corresponding data set(s). Dihedral angle mean absolute error (MAE) is computed analogously. Dihedral accuracy is defined as the fraction of dihedral angles with absolute difference less than 20° from the corresponding native angle. We further divide our results based on residue centrality. Core residues are defined as those amino acids with at least 20 *C_β_* atoms within a 10Å radius of the target residue’s *C_β_* atom. surface residues are defined as those amino acids with as most 15 *C_β_* atoms within the same region of interest. All Comparisons are made on native structures from CASP13, and CASP14 (see Table S11 for a list of targets in each test set).

We begin by comparing the average RMSD over CASP13 and CASP14 test sets (see Table S6 for results on CASP13-FM and CASP14-FM test sets). Table 1 shows that variants of our methods consistently achieve the lowest RMSD in each centrality category across all data sets. The inclusion of triangle updates yields a significant improvement over the TFN-Transformer alone. Compared to the other deep learning method DLPacker, our method obtains much lower RMSD in all categories, and observes the largest improvement on surface residues. Interestingly, the impact of triangle updates is less substantial on protein surface residues suggesting that triangle updates play a larger role in determining conformations protein core residues. We explore this further in Figure 5.

**Table 1:** Average RMSD (Å) on the CASP13 and CASP14 targets. Results are divided by residue centrality (All, Core, and Surface).

| Method | RMSD (Å) |  |  |  |  |  |
| --- | --- | --- | --- | --- | --- | --- |
|  | CASP13 |  |  | CASP14 |  |  |
|  | All | Core | Surface | All | Core | Surface |
| SCWRL | 0.771 | 0.471 | 0.983 | 0.910 | 0.556 | 1.125 |
| FASPR | 0.751 | 0.481 | 0.960 | 0.899 | 0.564 | 1.112 |
| RosettaPacker | 0.713 | 0.395 | 0.948 | 0.864 | 0.473 | 1.087 |
| DLPacker | 0.633 | 0.408 | 0.815 | 0.797 | 0.499 | 0.994 |
| TFN-Transformer | 0.608 | 0.440 | 0.746 | 0.758 | 0.535 | 0.905 |
| +Tri | **0.550** | **0.347** | **0.707** | **0.697** | **0.432** | **0.874** |
| +Tri+Local | <u>0.557</u> | <u>0.354</u> | <u>0.715</u> | <u>0.701</u> | <u>0.438</u> | <u>0.875</u> |
| Residue Count | 18139 | 6443 | 7520 | 13399 | 4021 | 6107 |

In terms of residue-level RMSD, the most significant improvements are achieved for Arg and His, each of which have large positively charged side-chains, as well as bulky hydrophobic amino acids Phe, Trp, and Tyr. A full overview of residue-level RMSD on the CASP13 and CASP14 data sets can be found in Supplementary Tables S8, and S9.

As shown in Figure 4, adding triangle updates before the TFN-Transformer improves the overall average RMSD of Arg, His, Phe, Tyr, and Trp by 11%, 16%, 20%, 28%, and 26% respectively. For surface residues, the performance difference between the TFN-Transformer and TFN+Tri is significantly smaller. On the other hand, the performance gap between our method and others increases for surface residues. We outperform the next best method by 16%, 17%, 25%, 20%, and 28% respectively.

**Figure 4:**
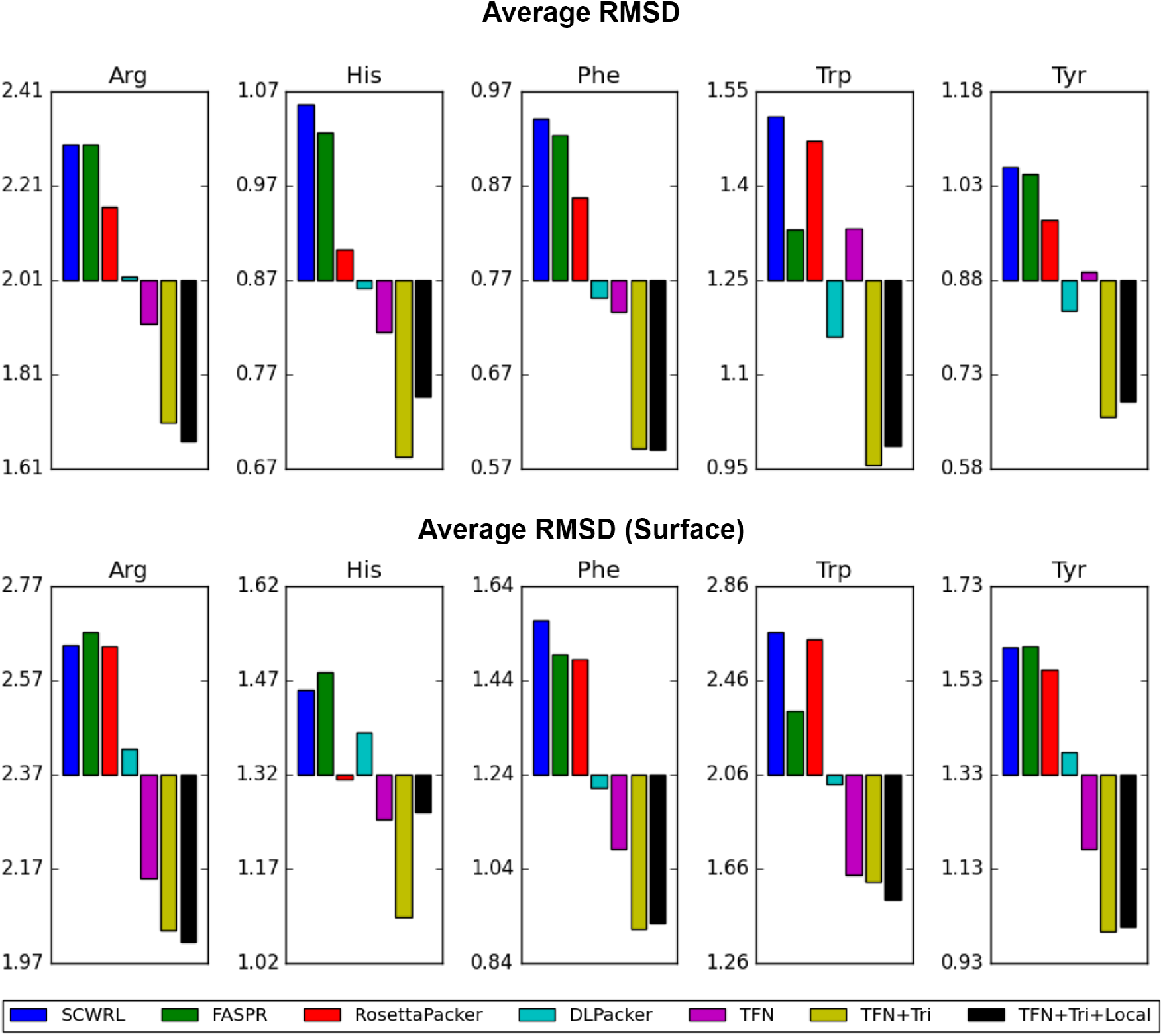
Average RMSD (Å) (y-axis) of each method (x-axis) on Arg, His, Phe, Trp, and Tyr. The RMSD values are averaged over the entire CASP14 data set for each residue type. Average RMSDs for instances of each residue type appearing on the protein’s surface are shown in the bottom plot.

We further investigated the role of centrality in side-chain reconstruction accuracy by measuring average RMSD and average χ_1–4_ MAE with respect to the number of *C_β_* atoms in a residue’s microenvironment. Not surprisingly, RMSD and chi angle prediction error decrease rapidly as centrality increases. In Figure 5, we verify that the marginal improvement of triangle updates increases with centrality. For both RMSD and MAE, the performance gap between TFN and TFN+Tri, increases with centrality, suggesting that triangle updates are important for accurately determining side-chain conformations in protein cores. The opposite is true when comparing TFN+Tri with DLPacker, where the gap decreases as centrality increases.This is not surprising, as DLPacker iteratively constructs each residue’s side-chain using only atoms the residue’s immediate microenvironment as input. This choice of input features implies that features for protein surface residues are more sparse than those for core residues.

**Figure 5:**
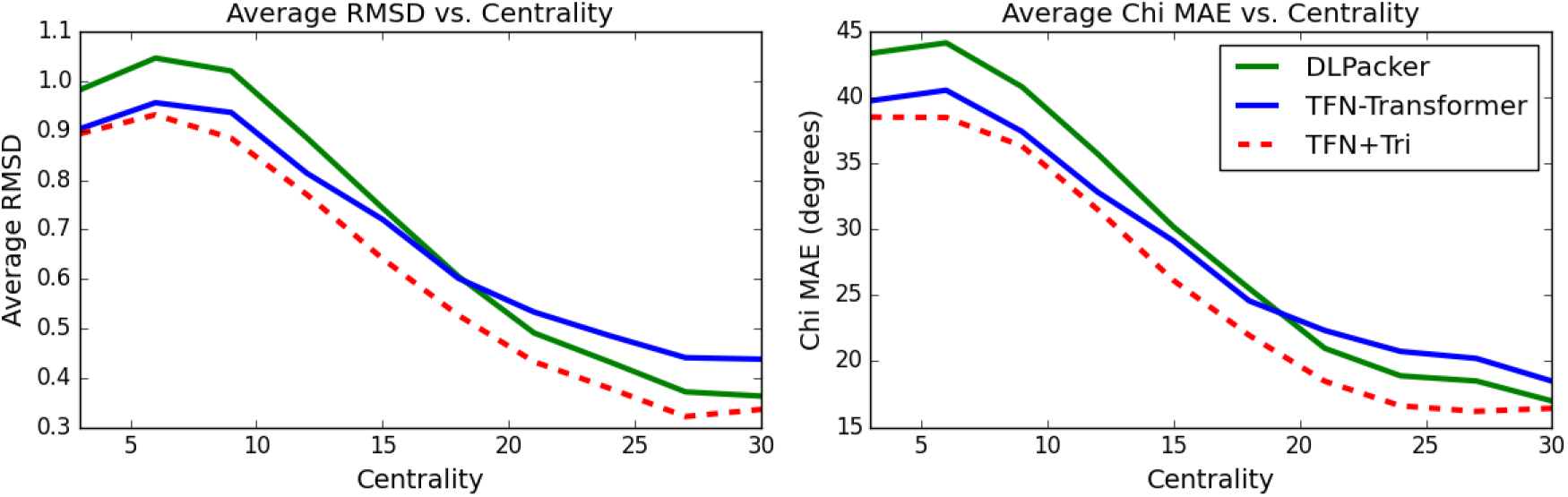
Plots of average RMSD and χ_1–4_-MAE° against centrality for methods DLPacker, TFN-Transformer (TFN), and TFN-Transformer with Triangle Updates (TFN+Tri). Values are computed using all targets in both CASP13 and CASP14 data sets. Values were disregarded if the number of residues with the corresponding centrality was less than 30.

In addition to achieving top performance in reconstructed side-chain RMSD, our method is also significantly faster than all other methods except for FASPR. Table 2 shows the cumulative and relative time spent by each method for reconstructing side-chains of all targets in the CASP13 dataset.

**Table 2:** Time comparison of PSCP methods. Relative times for reconstructing the side-chain atoms of all 83 targets in the CASP13 dataset. Run on a single RTXA6000 GPU, our method with local triangle updates is able to reconstruct all side-chain atoms in 68 seconds.

| Method | Ours | DLPack | Ros. Pack | FASPR | SCWRL4 |
| --- | --- | --- | --- | --- | --- |
| **Cumulative Time (s)** | 68.3 | 8499.6 | 10368.2 | 32.5 | 1001.9 |
| **Relative Time** | 1.0 | 124.4 | 151.7 | 0.5 | 14.7 |

We now turn our attention to dihedral angle prediction accuracy. As pointed out by Zhang et al.[15], the prediction accuracy of side chain dihedrals is much sharper when all chi dihedrals are considered, and a 20° cutoff is used when comparing to the native structure. We opt to use this criteria for all reported accuracy. In addition, we assess the quality of each method in terms of χ_1–4_ MAE with the native structure.

We analyzed side-chain dihedral accuracy on the CASP13 and CASP14 data sets and show the results in Table 3. In terms of χ_1–4_ MAE, the TFN transformer with triangle updates achieves top-1 or top-2 performance in each data set. The performance carries over into prediction accuracy, where the method also achieves top 1 or top 2 performance on each data set regardless of residue centrality. Local triangle updates achieve competitive performance, with top-2 MAE scores for all but one category.

**Table 3:** Side-chain dihedral prediction results on the CASP13 and CASP14 targets. Accuracy results shown for all, core, and surface residues.

|  | MAE<sup>o</sup> |  |  |  | Accuracy |  |  |
| --- | --- | --- | --- | --- | --- | --- | --- |
| | $\chi_1$ | $\chi_2$ | $\chi_3$ | $\chi_4$ | All | Core | Surface |
| CASP13 |  |  |  |  |  |  |  |
| SCWRL | 25.89 | 26.18 | 46.55 | 53.79 | 67.6% | 59.3% | 81.3% |
| FASPR | 25.22 | 25.36 | 47.25 | 52.93 | 68.1% | 59.9% | 80.3% |
| RosettaPacker | 24.15 | 24.49 | <u>44.84</u> | 52.19 | 69.4% | 60.6% | **83.7%** |
| DLPacker | 20.63 | 23.39 | 46.48 | 66.10 | 69.0% | 61.1% | 81.3% |
| TFN-Transformer | 19.87 | 23.03 | 46.71 | 54.42 | 68.6% | 62.0% | 78.7% |
| +Tri | **18.00** | <u>21.15</u> | **43.58** | <u>52.03</u> | **71.2%** | **63.3%** | <u>82.9%</u> |
| +Tri+Local | <u>18.69</u> | **20.95** | 45.07 | **51.63** | <u>70.4%</u> | <u>62.5%</u> | 82.3% |
| CASP14 |  |  |  |  |  |  |  |
| SCWRL | 30.92 | 29.63 | 49.07 | 51.24 | 61.3% | 74.9% | 54.4% |
| FASPR | 30.55 | 28.88 | 47.91 | 52.54 | 62.0% | 74.0% | 55.8% |
| RosettaPacker | 29.33 | 28.94 | 47.19 | 52.09 | 62.7% | **77.7%** | 55.5% |
| DLPacker | 26.65 | 27.49 | 51.58 | 68.13 | 61.8% | 74.9% | 54.2% |
| TFN-Transformer | 25.95 | 27.00 | 48.53 | 51.96 | 61.9% | 72.6% | <u>56.0%</u> |
| +Tri | **24.06** | **24.81** | **45.42** | **50.79** | **65.1%** | <u>77.4%</u> | **57.8%** |
| +Tri+Local | <u>24.44</u> | <u>25.42</u> | <u>46.29</u> | <u>50.84</u> | <u>63.4%</u> | 76.4% | 55.9% |

In line with the results reported by Misiura et al.[21], when compared to traditional methods, deep learning methods recover *χ*_1_ dihedral angles considerably closer to those of the native structure. For our methods, this improvement carries over to *χ*_2_ angle prediction, where triangle updates obtain an 8% and 10% improvement over the next best method DLPacker on the CASP13 and CASP14 targets.

To better understand the instances where our method loses its advantage to traditional PSCP algorithms, we consider the performance on four amino acids with high side-chain dihedral degrees of freedom. Following DLPacker, we consider dihedral prediction for charged and polar amino acids Lys, Arg, Glu, and Gln in Table 4. For these amino acids, variants of our methods are still competitive with traditional PSCP algorithms, comparing favorably in *χ*_1–2_ MAE but losing their edge at higher orders. Comparing accuracy scores tells a different story. Although we are able to obtain comparable or lower MAE values for each degree of freedom, we only obtain top-1 or top-2 accuracy for Gln and Glu. On the other hand, physics-based RosettaPacker obtains top-1 or top-2 performance for each amino acid along with top-1 *χ*_4_ scores. As the authors of DLPacker also point out, new training methods or loss functions which improve on higher order dihedral accuracy is an important area for future research.

**Table 4:** Dihedral MAE on the CASP13 targets for charged and polar Amino Acids with high degrees of freedom.

|  | MAE° |  |  |  |  | MAE° |  |  |  |
| --- | --- | --- | --- | --- | --- | --- | --- | --- | --- |
| | $\chi_1$ | $\chi_2$ | $\chi_3$ | $\chi_4$ | acc. | $\chi_1$ | $\chi_2$ | $\chi_3$ | acc. |
|  | ARG |  |  |  |  | GLN |  |  |  |
| SCWRL | 31.4 | 25.3 | 60.1 | 55.7 | <u>0.550</u> | 33.8 | 45.2 | 35.3 | 0.559 |
| FASPR | 34.1 | 25.9 | 63.2 | 57.5 | 0.537 | 33.4 | 42.5 | 36.1 | 0.558 |
| RosettaPacker | 31.1 | <u>24.3</u> | 60.6 | **51.4** | **0.564** | 29.3 | 36.6 | 35.5 | **0.591** |
| DLPacker | 25.6 | 26.2 | <u>57.9</u> | 61.1 | 0.516 | 31.2 | 39.3 | 34.7 | 0.576 |
| TFN-Transformer | 25.0 | 24.9 | 62.0 | 58.3 | 0.475 | <u>27.7</u> | 33.5 | 35.8 | 0.558 |
| +Tri | **21.8** | **22.1** | **56.9** | 52.8 | 0.522 | **25.2** | <u>29.3</u> | <u>34.0</u> | <u>0.587</u> |
| +Tri+Local | <u>22.5</u> | <u>24.3</u> | 58.8 | <u>52.5</u> | 0.491 | <u>27.7</u> | **28.4** | **32.9** | 0.580 |
|  | LYS |  |  |  |  | GLU |  |  |  |
| | $\chi_1$ | $\chi_2$ | $\chi_3$ | $\chi_4$ | acc. | $\chi_1$ | $\chi_2$ | $\chi_3$ | acc. |
|  | LYS |  |  |  |  | GLU |  |  |  |
| SCWRL | 28.6 | 31.9 | <u>40.3</u> | <u>48.9</u> | 0.617 | 38.1 | 36.6 | 36.1 | 0.540 |
| FASPR | 28.0 | **30.1** | 42.8 | 49.5 | **0.629** | 35.8 | 38.3 | 36.6 | 0.542 |
| RosettaPacker | 27.0 | 31.7 | **40.2** | **48.5** | <u>0.623</u> | 36.1 | 37.3 | 35.7 | <u>0.547</u> |
| DLPacker | 22.7 | 30.4 | 47.8 | 71.0 | 0.522 | 30.5 | 37.5 | <u>34.6</u> | 0.532 |
| TFN-Transformer | 21.7 | 32.1 | 42.3 | 50.5 | 0.577 | 28.5 | 35.41 | 32.3 | 0.546 |
| +Tri | **18.6** | 30.3 | 40.4 | 50.5 | 0.594 | **25.9** | **32.9** | **31.0** | **0.583** |
| +Tri+Local | <u>19.8</u> | **30.1** | <u>40.3</u> | 50.0 | 0.595 | <u>26.7</u> | <u>34.1</u> | 35.6 | <u>0.547</u> |

### 3.1 Ablation and Architecture Assessment

We consider several variants of SE(*k*)-equivariant self attention. Each variant can be categorized by the operation used to compute the similarity between keys and queries of different features types (i.e. 1D-scalar and 3D-point features) and the operation used to compute the final attention weights. We focus on the well-known dot product similarity, used in the SE(3)-Transformer and negative distance similarity which was first presented in the Invariant Point Attention module of AlphaFold2. Aside from calculating similarity between points, these architectures also differ in their calculation of attention scores - AlphaFold2 uses the same shared attention weights for scalar and point features, whereas the SE(3)-Transformer computes attention weights for each type. This is outlined more formally in Table 5. Following the conventions of ([27, 12]), we use a superscript ℓ to denote type-ℓ features of dimension 2ℓ +1.

**Table 5:**
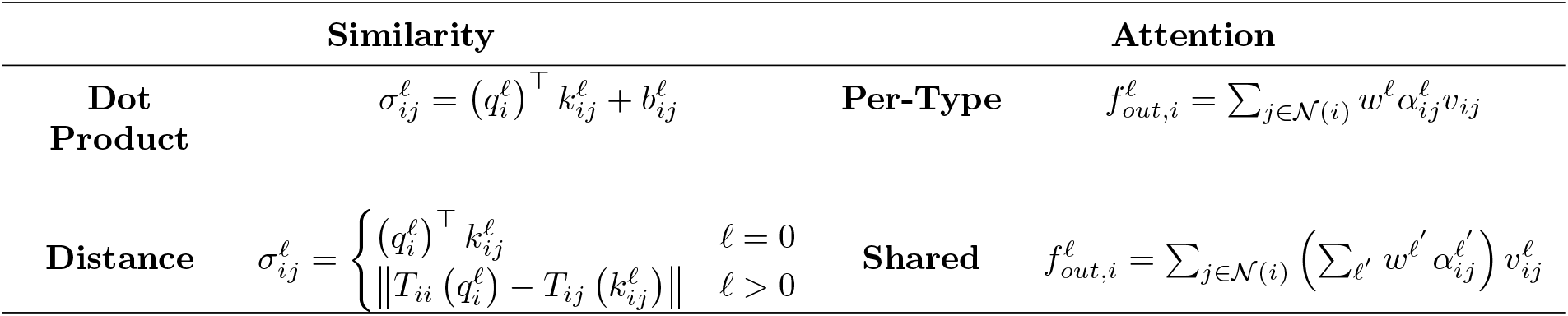
Description of similarity and attention types. Here, we use a superscript ℓ to denote type-ℓ features of dimension 2ℓ +1 (i.e. ℓ = 0 for scalar features and ℓ =1 for point features). For distance similarity, we use *T_xy_* to denote some transformation mapping points into so-called “local frames” of the respective node used to ensure equivariance of the operation.

We trained four TFN-Transformer models differing only by attention and similarity type. Each model used a hidden dimension of 24 for point features and 180 for scalar features. Ten attention heads were used in each of eight attention blocks. A head dimension of 20 was used for scalar features and 4 was used for point features. A maximum of 14 nearest neighbors were considered and a radius of 16Å was used as a cutoff. The average RMSD and dihedral accuracy results of the four attention variants are shown in Table 6.

**Table 6:**
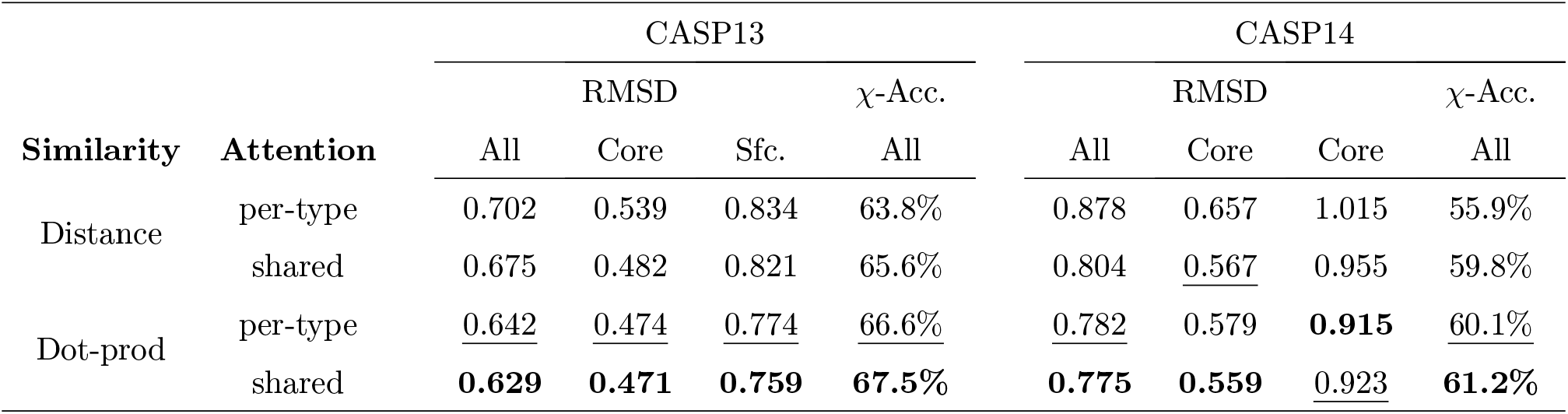
Comparison of attention and similarity operations on performance. Performance is measured using average residue RMSD (Å) and χ_1–4_ prediction accuracy.

The results in Table 6 show that shared attention weights produce better results for both similarity types. Overall, dot-product-based similarity outperforms distance based similarity, even when per-type attention is used. As pointed out by Fuchs et al[12], this may be due to the fact that each basis kernel in a TFN is completely constrained in the angular direction. By using dot-product based similarity, the angular profile of the basis kernels are modulated by the attention weights.

We also experimented with different architectural variants. First, we used a linear projection, rather than a TFN to compute keys at each attention head. Next, we augmented the input to the TFN radial kernel by concatenating the pairwise distances between hidden coordinates. Third, we tried removing the attention calculation between points and instead used weights derived from scalar features for pointwise attention. With each variant, shared attention and dot product attention was used. As shown in Table 7, using a linear projection for attention keys has the largest impact on RMSD score - surprisingly more than removing point-based attention all together. This suggests that TFN neighbor convolutions are an important component of the transformer architecture. The results also show that RMSD scores are improved when pairwise distances between hidden coordinate features are concatenated to the input of the TFN radial kernel.

**Table 7:**
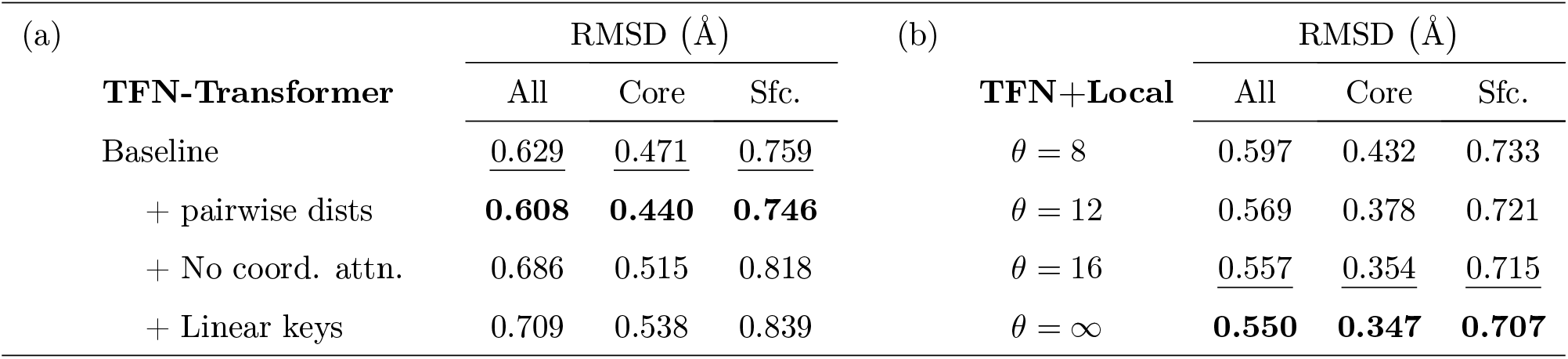
The tables show average side-chain RMSD on the CASP13 targets. Table (a) shows the effect of TFN architectural features. *Baseline* denotes our TFN-Transformer with shared dot-product attention and pair bias. The other three rows show results after changing some component. Table (b) shows the effect of neighbor distance threshold on RMSD with *θ* = 8, 12, 16 and ∞. Each model in (b) uses our locality aware graph transformer with distance cutoff *θ* and TFN-Transformer with shared dot-product similarity, pair bias, and pairwise distance features. All other hyperparameters were held constant.

| (a) | RMSD (Å) |  |  | (b) | RMSD (Å) |  |  |
| --- | --- | --- | --- | --- | --- | --- | --- |
| TFN-Transformer | All | Core | Sfc. | TFN+Local | All | Core | Sfc. |
| Baseline | <u>0.629</u> | <u>0.471</u> | <u>0.759</u> | $\theta = 8$ | 0.597 | 0.432 | 0.733 |
| + pairwise dists | **0.608** | **0.440** | **0.746** | $\theta = 12$ | 0.569 | 0.378 | 0.721 |
| + No coord. attn. | 0.686 | 0.515 | 0.818 | $\theta = 16$ | <u>0.557</u> | <u>0.354</u> | <u>0.715</u> |
| + Linear keys | 0.709 | 0.538 | 0.839 | $\theta = \infty$ | **0.550** | **0.347** | **0.707** |

We also considered the effect of the neighbor distance threshold on performance. This threshold is used as the maximum valid edge length for triangle updates, and to determine residue adjacency in the locality aware graph transformer. It is also used to define the neighborhood of scalar and point features in the TFN-Transformer. We tried three distance thresholds, 8Å, 12Å, and 16Å The results are shown in Table 7.

Average RMSD clearly decreases with increasing neighbor distance. The marginal improvement diminishes with increasing radius, and the bulk of the improvement comes from residues in the protein core.

## 4 Concluding Discussion

In this work, we developed AttnPacker, an SE(3)-equivariant model for direct prediction of side-chain coordinates based on backbone geometry. AttnPacker uses spatial information derived from protein backbone coordinates to efficiently model residue and pairwise neighborhoods. This, coupled with an SE(3)-equivariant architecture, allows for the simultaneous prediction of all amino acid side-chain coordinates without the use of rotamer libraries or conformational sampling.

Components of our model were inspired by AlphaFold2 and the SE(3)-Transformer. By generalizing and carefully evaluating ideas from these architectures we were able to achieve better efficiency or better than the original implementations. Specifically, we generalized the Evoformer module of AlphaFold2 to spatial graphs specific to protein backbones. We also modified the attention heads of the SE(3)-Transformer to incorporate pair-bias, distance information between hidden points, and shared dot-product similarity.

Our TFN-Transformer outperforms all other methods in terms of average RMSD on the CASP13 and CASP14 targets, even without triangle updates for pair-features. On the other hand, the baseline SE(3)-Transformer with per-type attention falls short of DLPacker for the CASP13 targets. Part of this improvement is achieved by augmenting edge features with pairwise distance information between hidden points. We hypothesize that this information is especially important for deeper TFN-based architectures. TFNs require a basis of spherical harmonics which is typically computed once on the relative positions of input points and then shared across all proceeding layers. As a result of using spherical basis functions, information about relative distances between hidden coordinates is lost. To account for this, distances between pairs of initial point features are typically concatenated to the input of the TFN radial kernel. Traditionally, this input is the same across all layers - the pairwise distance *d_ij_* between initial points *i* and *j*, and (optionally) a fixed vector of pair feature *e_ij_*. In theory, distance information between hidden point features could be captured by TFNs convolution operation, but we believed that incorporating this information at the input level could be beneficial. Since separate radial kernels are learned for each TFN, we chose to augment our input with distance information between hidden coordinate features at each layer, resulting in a modest improvement over the baseline implementation.

The difference in performance between our model trained using full triangle updates and our model trained with local triangle updates is very small, especially considering the fact that that all other hyperparameters were the same for each model. Restricting residue and pairwise attention updates to at most 30 nearest neighbors did not considerably degrade performance. On the other hand, this drastically reduces the memory required to train the model. In the future, it would be interesting to see if a larger or deeper model could improve performance further.

On top of outperforming other popular methods, our model presents several other advantages. First, it is extremely fast. We are able to predict all side-chain conformations for a 500-residue protein in less than a second using a single nvidia RTXA6000 GPU. On the other hand, DLPacker must be run iteratively for each amino acid side-chain causing a large dependence on protein length. OPUS-Rota4 also requires several pre- and post-processing steps in the form of derived constraints and gradient descent. Since our method directly predicts side-chain coordinates, the output is fully differentiable which benefits downstream prediction tasks such as refinement or protein-protein interaction. This also circumvents the use of engineered energy functions and rotamer libraries and places the emphasis on architectural innovations and better loss functions.

Our model is also very simple to use - it requires only a pdb file to run. In contrast, OPUS-Rota4 requires voxel representations of atomic environments derived from DLPacker, logits from trRosetta100, secondary structure, and constraint files derived from the output of OPUS-CM. Obtaining the requisite input data made it too difficult for us to compare results with this method.

In addition to our method’s speed and simplicity, it further succeeds in efficiently modeling residue-level local environments by using locality-based graph attention during feature and structure generation stages. On the other hand, DLPacker and OPUS-Rota4 use 3D-voxelized representations of each amino-acid’s microenvironment - requiring space *O*(*v*^3^*cd*), where *v* is the voxelized width (40 in the case of DLPacker and OPUS), c is the number of channels, and d is the channel dimension. Although this choice of representation has helped facilitate good performance for each method, the memory requirements prevent simultaneous modeling of all amino acid sidechains which could ultimately hinder reconstruction accuracy. We hypothesize that simultaneously modeling all side-chains helps contribute to our method’s RMSD improvements for surface residues.

Although AttnPacker yields significant improvements in residue-level RMSD, we remark that traditional PSCP methods SCWRL, FASPR, and RosettaPacker still perform comparably in terms of *χ*_3_ and *χ*_4_ angle prediction. We suspect that incorporating dihedral information into our model - either directly or through an appropriate loss function - could help improve performance.

## 6 Acknowledgements

M.M. would like to thank Phil Wang for some support code and discussions on SE(3)-equivariant architectures.

## 7 Author contributions

J.X. conceived and supervised the project and built the in-house training data. M.M. developed, implemented and tested the algorithm. M.M. and J.X. analyzed the results and wrote the manuscript.

## Supplementary Material

### S1 Data Collection

SCWRL4(ver 4.02), and FASPR were run with default configurations. Rosetta’s fixbb application was run with nonflexible backbone coordinates and the maximum number of rotamers *by passing -EX1, -EX2, -EX3* and *-EX4* flags. We also included flags *-packing:repack_only* to disable design, *-no_his_his_pairE*, and *-multi_cool_annealer 10* to set the number of annealing iterations - these settings are recommended in the rosetta tutorial. We ran Rosetta Packer 5 times for each target protein using Rosetta’s ref2015 energy function and selected the conformation with lowest energy. DLPacker was run using the [pre-trained release from the author's github](https://github.com/nekitmm/DLPacker), downloaded Sept. 17th, 2021. Side-chains were reconstructed in non-increasing order of the number of the number of atoms in the corresponding amino acid’s microenvironment.

We ran each method on each of the targets listed in Table S11. All of the methods were run using the native backbone coordinates as input. For our models, we considered only the first 800 amino acids and corresponding coordinates in each target’s PDB file. The output of FASPR, SCWRL, Rosetta Packer, and DLPacker was cropped to the same length where applicable.

### S2 Overview of Hyperparameters

We tried to maintain consistent hyperparameters for all models. We mainly tuned parameters for model depth, number of nearest neighbors, number of attention heads, head dimension, and the distance at which residues or pair features should be considered neighbors. We settled upon the hyperparameter values listed in Table S1. In choosing the parameters, we aimed to balance memory usage with model capacity in each submodule. The final settings required ^~^32GB of GPU memory during training when full triangle updates are used (this is based on a maximum sequence length of 300 residues). The actual memory usage is lower when local triangle updates are used.

**Table S1:** Hyperparameters used for each model (unless otherwise specified). A description of input feature dimensions can be found in Section S4.

|  | **Graph**<br>**Transformer**<br>Full Tri. Updates<br>(*residue, pair*) | **Graph**<br>**Transformer**<br>Local Tri. Updates<br>(*residue, pair*) | **TFN-Transformer**<br>(*scalar, point*) |
| --- | --- | --- | --- |
| Depth | 12 | 12 | 8 |
| Hidden Dim. | 180, 128 | 180, 128 | 128, 24 |
| Num. Attention<br>Heads | 10, 4 | 10, 4 | 10, 10 |
| Head Dim. | 20, 32 | 20, 32 | 20, 4 |
| Neighbor Distance<br>Cutoff | N/A | 15, 15 | 15 |
| Max. Nearest<br>Neighbors | N/A | 30, 30 | 14 |

### S3 Training Details

We trained and validated all models using the BC40 training data set (SOURCE) and validated on BC40 validation set. The data set contains ^~^36k proteins which are selected from PDB database by 40% sequence identity cutoff. We first train our models for 10 epochs with a sequence crop size of 300, and an initial learning rate of 10^−3^. The learning rate is decreased by a factor of two every three epochs, and we do not use any warm-up. After the first stage completes, the models are trained for an additional 2 epochs with a learning rate of 10^−4^ and a sequence crop size of 500. No other parameters are modified between the two training stages.

To optimize our models, we use Adam[19] with parameters *β*_1_ = 0.9, *β*_2_ = 0.999, and *ϵ* = 10^−8^, and use a minibatch size of 32. To stabilize training and avoid using learning rate warm-up schedules, we use ReZero[3], for every residual connection in our transformer blocks. We also apply gradient clipping by global norm[24] to clip the gradients of each example in a minibatch to have ℓ_2_ norm at most 1.

We apply gradient checkpointing on the triangle attention logits and TFN kernel outputs. This yields a massive decrease in memory consumption during training, at a cost of a ≈ 50% decrease in speed. Details are provided in Section S8. Overall, each model was trained for roughly six days on a single nvidia RTX A6000 gpu (^~^5 days for the first stage, and ^~^1 day for stage 2).

#### S3.1 Loss Function

Our loss function consists of two equally-weighted terms. The first is an auxiliary loss over predicted distances of distal side-chain atoms (see ’tip-atom’ defined in [13]). This term is applied to the pair output of the locality aware graph transformer. The pair output is first symmetrized, and then logits are obtained by linearly projecting into 46 bins covering 2Å – 20Å. Two bins are also added for predicted distance less than 2Å and greater than 20Å. If locality aware attention is used, the loss is only computed for pair ij if the corresponding residues are adjacent. Pairwise distograms are obtained by taking a softmax of the logits, and an averaged cross entropy loss is then applied.

The second loss term is applied to the predicted coordinates. Let 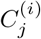 denote the *j^th^* side-chain atom coordinate for residue *i* in the native structure. Define 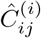 analogously for the predicted structure. The loss is computed as

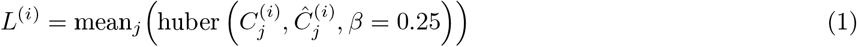

Where huber(*x, y, β*) is the Huber loss between *x* and *y* with smoothing parameter *β*. The final loss is computed as mean_*i*_ (*L*^(*i*)^).

Some care must be taken in computing Equation (1), since some residues have symmetric sidechains. For these residues, we consider all possible symmetries by swapping the coordinates of symmetric atoms and take the lesser of the swapped and not-swapped loss for the respective residue. A list of residues with symmetric sidechains and pairs of atoms for which we swap coordinates can be found in Table S2.

**Table S2:**
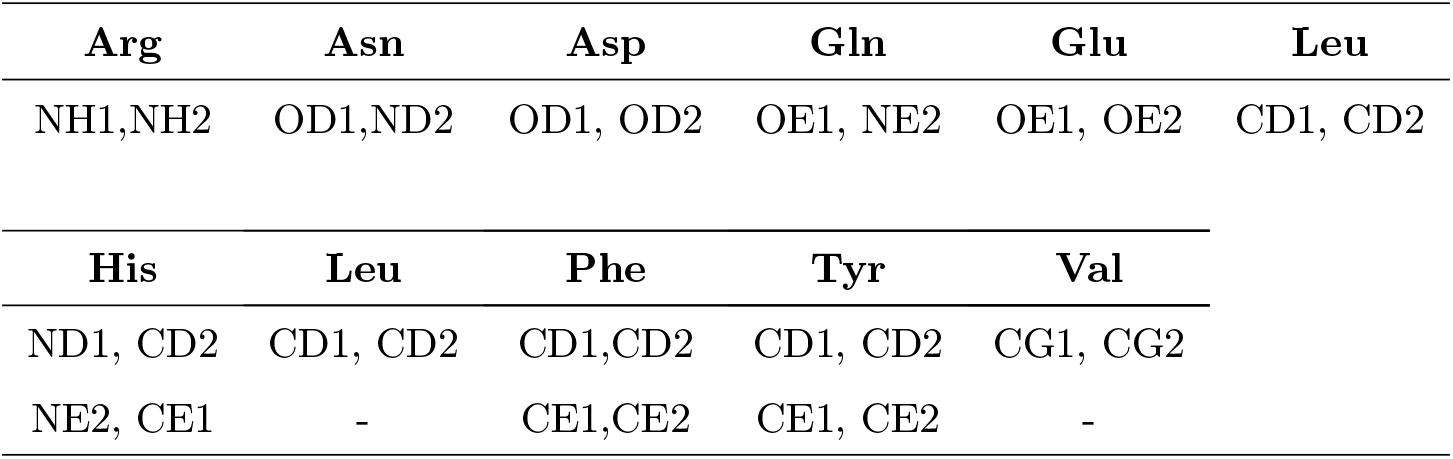
Amino acids with sidechain symmetries and the atom pairs which constitue these symmetries.

| Arg | Asn | Asp | Gln | Glu | Leu |
| --- | --- | --- | --- | --- | --- |
| NH1,NH2 | OD1,ND2 | OD1, OD2 | OE1, NE2 | OE1, OE2 | CD1, CD2 |
| His | Leu | Phe | Tyr | Val |  |
| ND1, CD2 | CD1, CD2 | CD1,CD2 | CD1, CD2 | CG1, CG2 |  |
| NE2, CE1 | - | CE1,CE2 | CE1, CE2 | - |  |

### S4 Model Input

#### S4.1 Input Features

Our model uses only input features derived directly from primary sequence and backbone coordinates. An overview of input feature types and the corresponding shape can be found in Table Table S3.

**Table S3:** Input features used by AttnPacker. The shape of the corresponding type for a protein with *L* residues is shown below each feature.

| Features Name & Shape | Description |
| --- | --- |
| res_type<br>[$L$] | A number in the range 0..21 representing the corresponding amino acid type |
| bb_dihedral<br>[$L, 3$] | Backbone phi, psi, and omega dihedral angles. |
| seq_pos<br>[$L$] | The sequence index of the respective residue, from 1..$L$ |
| centrality<br>[$L$] | The number of $C_\alpha$ atoms within 16Å of residue of the respective residue’s $C_\alpha$ atom. |
| atom_distance<br>[$L, L, 3$] | One-hot encoded binned distance between $C_\alpha - C_\alpha$, $C_\alpha - N$, and $N - O$ atoms in each residue. Each bin represents a distances from 2Å-20Å with two separate bins used for distances falling outside of this range. |
| tr_orientations<br>[$L, L, 3$] | Dihedral and planar angles defined by Yang et al[33]. |

We use standard residue level features and encodings for our input. These include a representation of amino acid type, binned relative sequence position, and embeddings/encodings of backbone dihedral angles. Less standard is the inclusion of residue centrality. The use of centrality based encodings was studied in [35], where the authors found that transformers significantly benefit from the addition of centrality encodings with graph-like data. We choose to incorporate this information at the input level, rather than in each attention block (as is proposed in [35]). Originally, we included SS3 secondary structure as part of our input, but found that this feature gave the same (if not slightly worse) results in terms of average residue RMSD scores.

For our pair embedding, we follow a scheme similar to [17], but also incorporate residue pair information following the approach of AlphaFold2. Rather than embed each residue pair separately, the authors propose using two separate embeddings for residue types. Denote the embeddings as *E_A_* and *E_B_*, then the pair feature for residues *i* and *j* of type *r_i_* and *r_j_* (resp.) is given by the outer-sum of *E_A_* (*r_i_*) and EB (*r_j_*). The authors further augment this information by add an embedding of the relative sequence separation between the two residues and we follow the same approach here.

**Figure S1:**
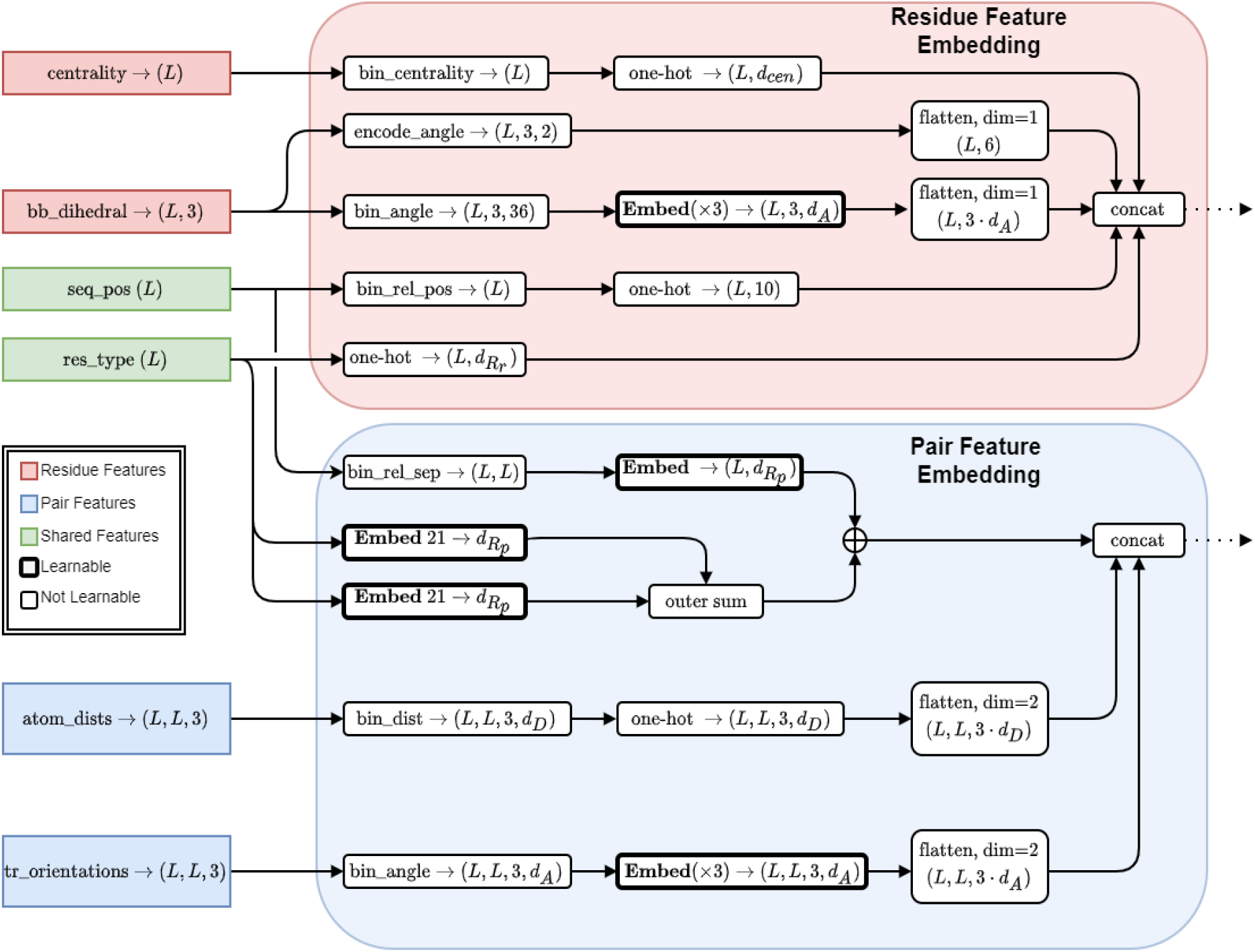
Overview of our input feature embedding. The feature shape is shown in parenthesis. Residue and pairwise features are embedded separately. Only information derived from primary sequence and backbone coordinates is considered. Full explanations of procedures referenced in this figure are given in Table S4. For residue features in our final models, we use *d_cen_ =* 7 for the one-hot encodings of centrality features, *d_A_* = 6 for our backbone torsion angle embedding dimension, and *d_R_r__ =* 32 for residue type embedding dimension. For pairwise features, we use *d_D_* = 34 for each one-hot distance encodings, *d_R_p__ =* 48 for residue pair embeddings, and *d_A_* = 6 for embeddings of trRosetta orientations.

**Table S4:** Description of Input Embedding Procedures referenced in Figure S1. The name of the procedure (top) along with the domain and range (bottom) are given in the first column. Here, we use *L* to denote the protein length.

| Procedure | Description |
| --- | --- |
| $\text{bin\_centrality}(c) :$<br>$\mathbb{N} \rightarrow [0..6]$ | Mapping given by $c \mapsto \min(\lfloor \frac{c}{8} \rfloor, 6)$ |
| $\text{bin\_angle}(\theta) :$<br>$(-\pi, \pi) \rightarrow [0..35]$ | Mapping given by $\theta \mapsto \lfloor 36 \cdot \frac{\theta + \pi}{2\pi} \rfloor$ |
| $\text{encode\_angle}(\theta) :$<br>$(-\pi, \pi) \rightarrow [0, 1]^2$ | Mapping given by $\theta \mapsto (\cos \theta, \sin \theta)$ |
| $\text{bin\_rel\_pos}(p) :$<br>$[1..L] \rightarrow [0..9]$ | Mapping given by $p \mapsto \lfloor 10 \cdot \frac{p-1}{L} \rfloor$ |
| $\text{bin\_rel\_sep}(s) :$<br>$[0..L-1] \rightarrow [0..48]$ | Maps the (signed) sequence separation $s$ to the index of a predefined interval. We chose intervals $[0, 1), [1, 2), [2, 3), [3, 4), [4, 5), [5, 6), [6, 7), [7, 8), [8, 10), [10, 12), [12, 15), [15, 20), [20, 30), [30, \infty)$ plus the negation of each interval. |
| $\text{bin\_dist}(d) :$<br>$\mathbb{R}^{\geq 0} \rightarrow [0..33]$ | Maps the pairwise distance $d$ to one of 32 equal width bins in the range $3\text{\AA}..20\text{\AA}$. Distances less than $3\text{\AA}$ and distances greater than $20\text{\AA}$ are placed into separate bins. |

### S5 Locality Aware Triangle Attention

Consider a set of points 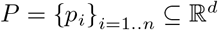. Define

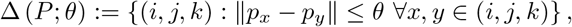

the set of triangles in *P* with maximum side length at most *θ*.

We develop a hypergraph approach to performing triangle updates 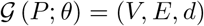, where

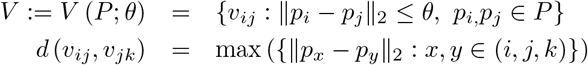

The edge set *E* is defined by the adjacency relation

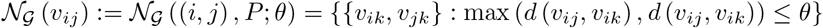

Observe that there is a node for every pair of points within distance *θ*, and the distance between two nodes sharing a common underlying point is the maximum edge length on the corresponding triangle.

When the underlying point set consists of backbone atom coordinates for a protein, a reasonable choice of *θ* results in a relatively small set of triangles. Furthermore, in order to efficiently compute triangle updates for each node *v_ij_*, we restrict 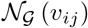 to the *N_e_* nearest neighbors under *d*(·, ·). This yields a 3-uniform, *O*(*N_e_*)- regular hypergraph with as many nodes as there are underlying points within distance *θ*. We denote the latter quantity by *N_v_*.

In theory, the pairwise features selected for triangle attention can be further restricted to the (at most) *N* nearest neighbors of each residue, as only the pair features for each residue’s *N* nearest neighbors are used to bias residue attention logits. In practice, we restrict pair features to the (at most) 2*N* nearest neighbors of each residue. More formally, the vertices of our triangle hypergraph consist of pairs (*i, j*) such that *j* is among the top 2*N* nearest neighbors of *i*. Triangle multiplication and attention updates then require time and space proportional to 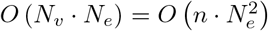 in total. These updates are identical to that of AlphaFold2, but restricted to this subset of triangles.

**Figure S3:**
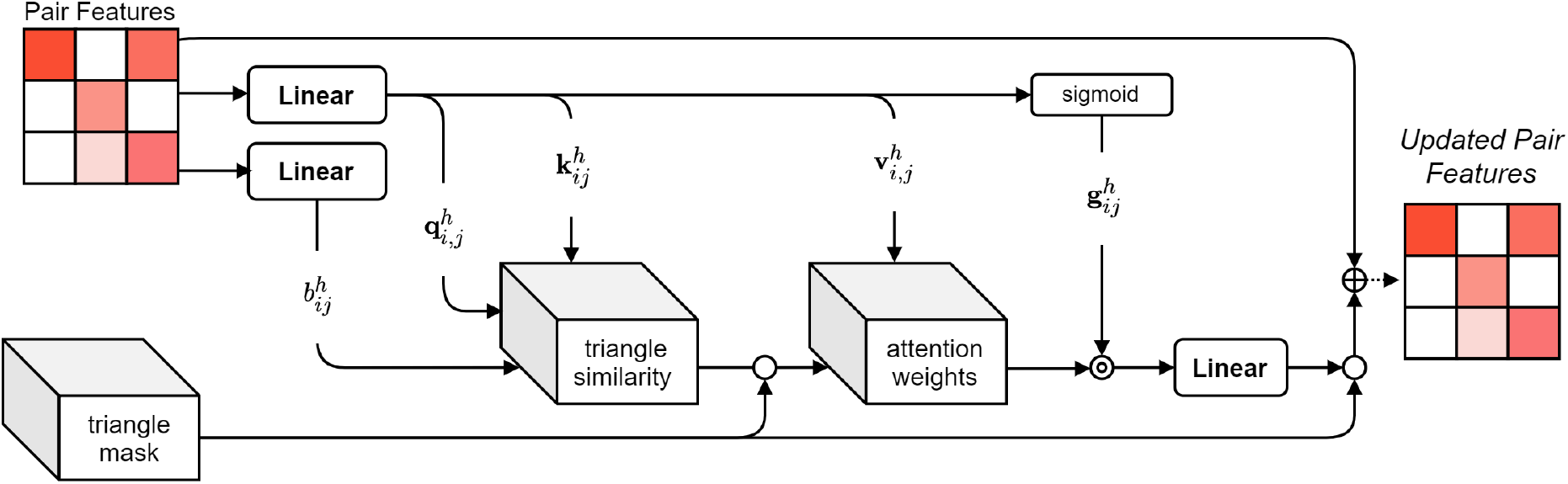
A simple implementation of local triangle attention updates. A mask of dimension *L* × *L* × *L* is used to control which triangles (and respective edge features) are updated during attention calculations. Masked positions are kept unchanged, and unmasked positions are filled with a large negative value (–10^6^) before softmax is applied to compute attention weights.

As an alternative, we can easily implement local triangle updates by computing a global triangle mask for each sample (see Figure S3). This approach does not yield the time and space savings previously discussed, but does serve as a simple drop-in replacement for existing models.

### S6 TFN-Transformer

#### Algorithm S1 SE(3)-Equivariant Normalization

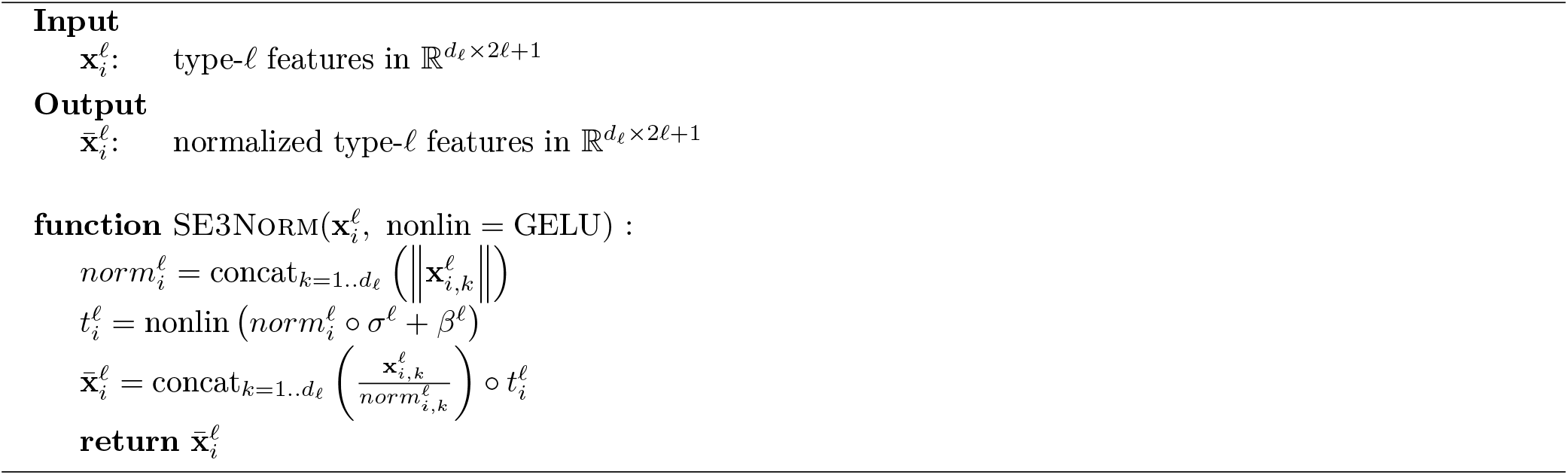

For normalization, we propose a method similar to Layer normalization with a norm-based non-linearity. Several papers have proposed SE(3)-Equivariant Normalization schemes (e.g. [12, 7, 16]), most include some form of Layer normalization [2], or restriction on the ℓ_2_ norm of coordinate features. In our experiments, we found that applying layer normalization to coordinate norms (and subsequently scaling by these values) sometimes caused instability in the early stages of training. In light of this, In Algorithm S1, we simply learn a scale and bias *σ*^ℓ^ and *β*^ℓ^ for each feature type.

#### Algorithm S2 Augment Edge Features

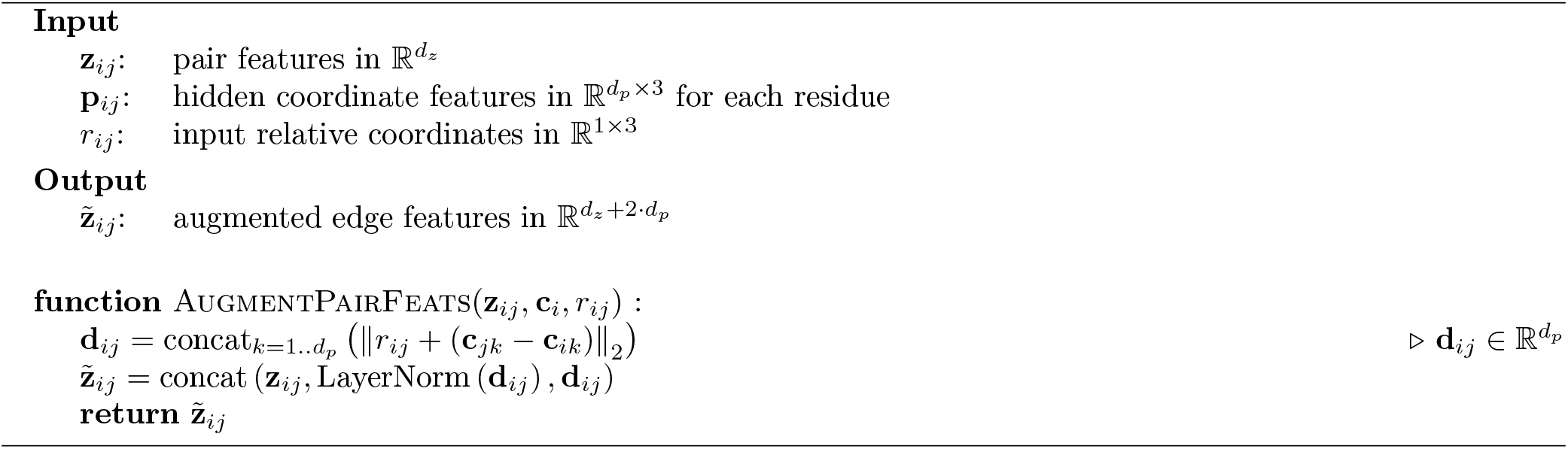

In augmenting the pair features with distance information, we choose to append both normalized and un-normalized distances between the hidden points. The output of this function is passed directly to the TFN radial kernel, which employs a 3-layer MLP with GELU nonlinearity to produce pairwise kernels for each pair of input feature types.

#### Algorithm S3 TFN-Transformer Attention Head

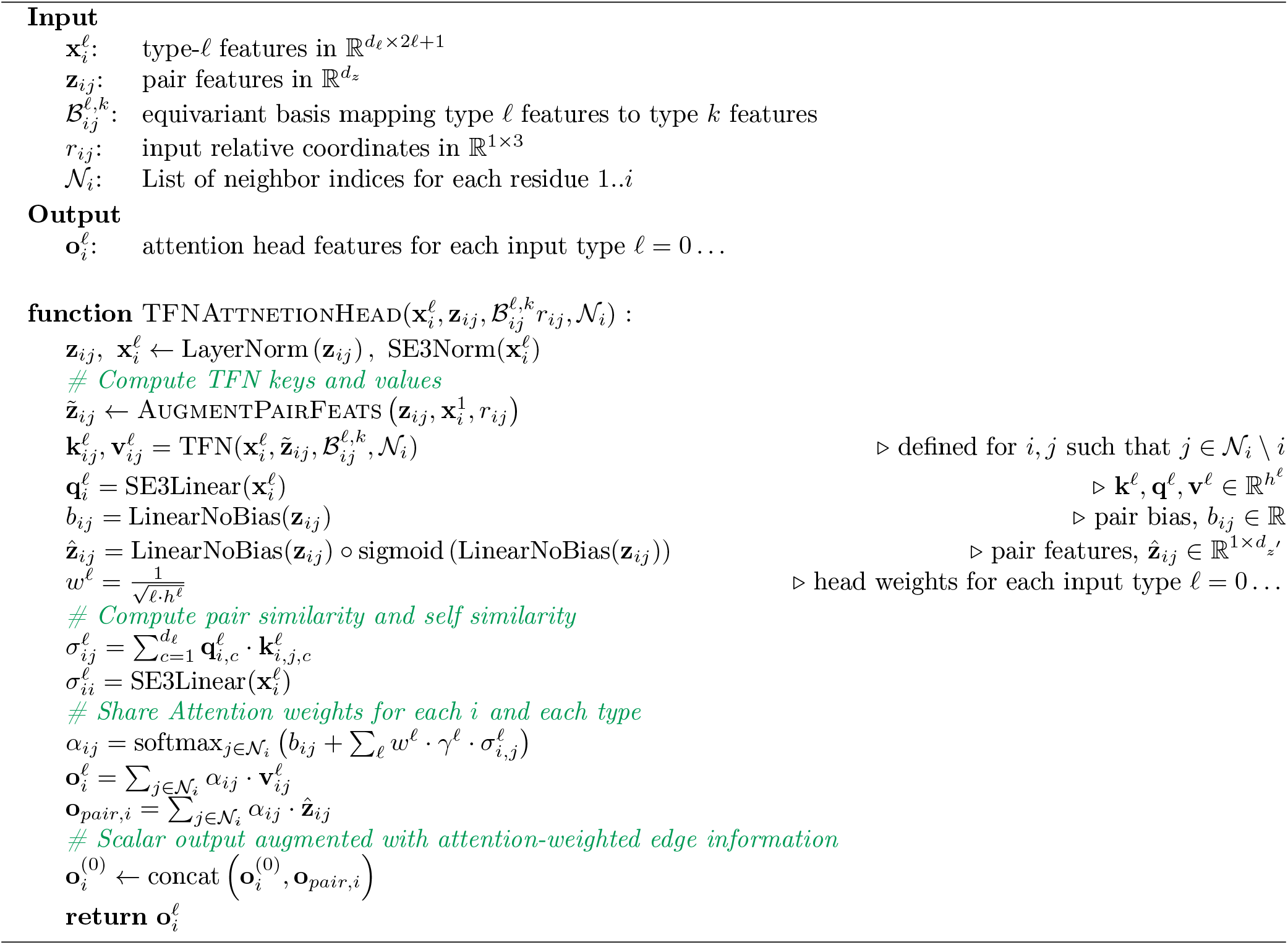

In Algorithm S3, we use *d*_ℓ_ to denote the input dimension, and to denote the hidden dimension (head dimension) of input feature type ℓ. Each attention head has a separate learnable weight for each input type ℓ. This weight is the softplus of a learnable scalar and is initialized so that *γ*ℓ = 1. For each input type, the output of each attention head is concatenated and linearly projected so that the output dimension matches the original input dimension. All linear projections (SE3Linear) follow the scheme proposed in [7].

We give a full description of out TFN-Transformer in Algorithm S4. Our TFN-Transformer consists of three components: (1) An equivariant mapping of scalar and coordinate input features to hidden feature types/dimensions (2) multiple TFN-based attention layers and (3) An equivariant mapping from hidden feature types/dimensions to output feature types/dimensions.

#### Algorithm S4 TFN-Transformer

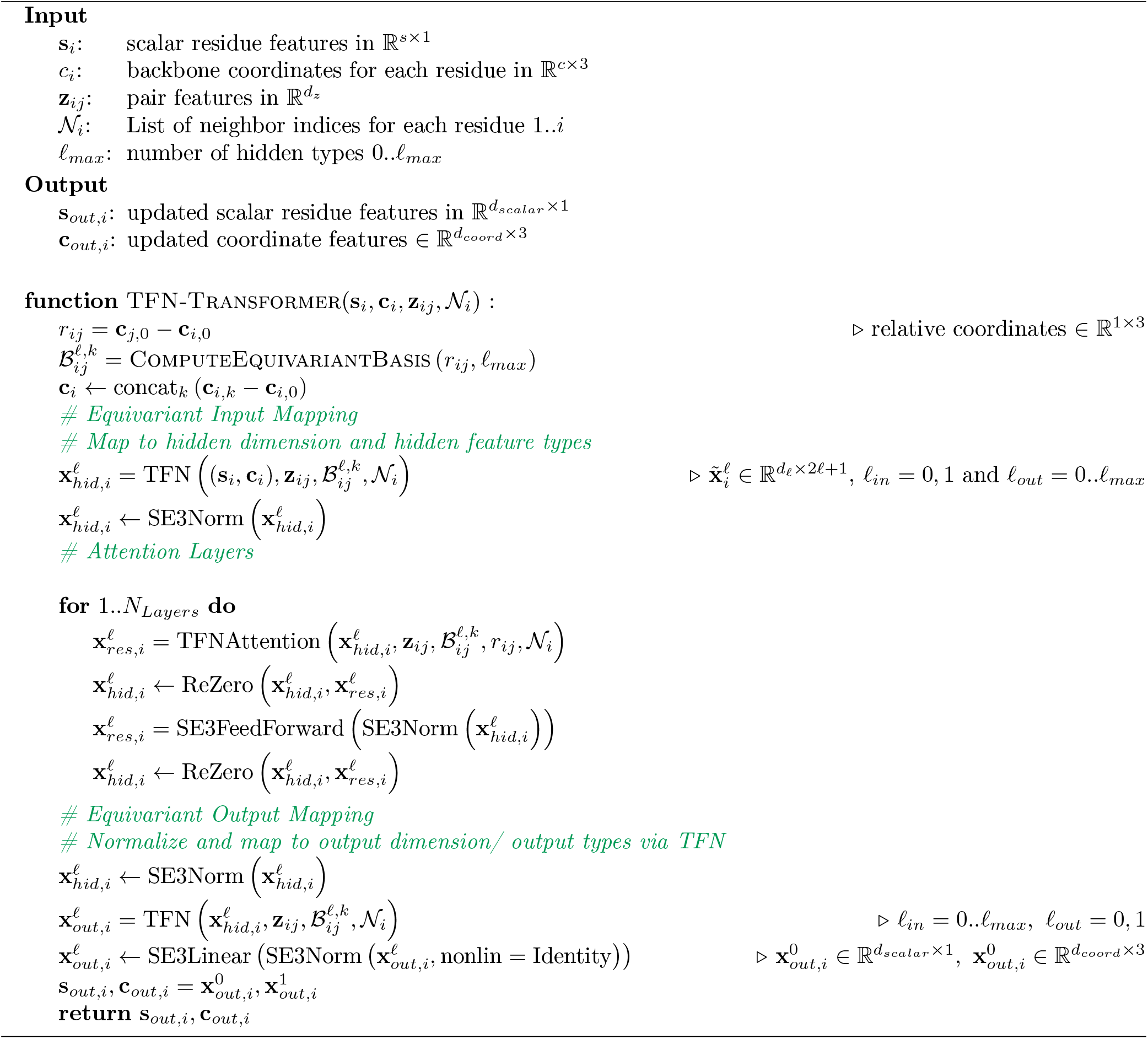

### S7 Full Architecture

**Figure S4:**
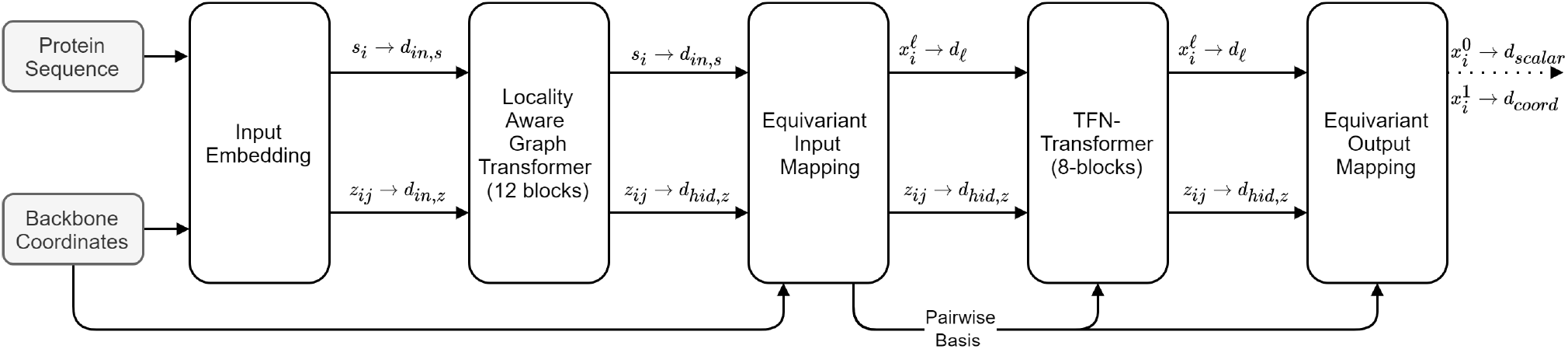
The full architecture of AttnPacker. we use *s_i_* to denote scalar (residue) features, and *z_ij_* to denote pair features. Backbone coordinates are used to define residue and pair adjacencies, and included in the equivariant input mapping (see Algorithm S4). The Equivariant input mapping returns type ℓ features 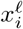 for user-defined types ℓ > 0. In our model, scalar output features 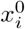 are discarded, and coordinate output 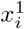 has *d_coord_* = 32, one channel for each possible side-chain atom type.

Our full architecture is shown in Figure S4. It consists of an input embedding (Figure S1) to produce scalar features for residues and pairs, and is followed by our locality-aware graph transformer and TFN-Transformer. Pair features are only modified in the locality-aware graph transformer block. The TFN-Transformer still makes use of pair features to produce radial kernels, bias attention logits, and augment the output of each attention head.

### S8 Memory Consumption

#### S8.1 Triangle Attention

In Section 2.2, we determine that the number of pair features requiring triangle updates is bounded by 2*NL*, where *N* is the maximum number of neighbors per point, and *L* is the protein length. Locality aware triangle attention stores at most *N* attention logits per pairwise feature. It follows that the number of attention logits stored for an attention layer with *h* heads is 2*N*^2^*L* · *h*. In practice, computing neighbor-wise attention can be costly. Depending on the implementation, repeating or grouping of neighbor features may cause the gradients for attention logits to require significantly more space. We experimented with several implementation, but ultimately decided to mask full triangle attention logits (*O*(*L*^3^) space) as a proof of concept. Masking attention logits effectively stops the gradient flow for all pair pair features which are not part of the triangle graph. This resulted in a modest improvement in memory, but more work will be needed to realize the full time and space benefits of this architecture.

**Table S5:** A comparison of memory usage for storing pair attention logits. *L* is the length of the input sequence. The third and fourth columns show the memory usage for an input sequence of length 300 and length 500 respectively. Values are calculated by fixing the number of heads *h* = 4, the depth *D* = 12, the number of nearest neighbors *N* = 30 per pair.

| | bits for attention logits<br>using 32-bit float<br>precision | $L = 300$ | $L = 500$ |
| --- | --- | --- | --- |
| Triangle Attention | $32 \cdot L^3 \cdot h \cdot D$ | 5.2 Gb | 24.0 Gb |
| Locality Aware<br>Triangle Attention | $32 \cdot 2N^2L \cdot h \cdot D$ | 0.10 Gb | 0.17 Gb |

#### S8.2 TFN Transformer

#### S8.3 TFN Memory Analysis

One of the main drawbacks of the SE(3)-Transformer is high memory usage caused by computing equivariant pairwise kernels in each attention block. To alleviate some of this overhead, we modify the TFN implementation used in the original SE3-Transformer proposed by Fuchs et al.[12]. Given input feature tensors of size (*d_in_*, *o_in_*), (*d_out_*, *o_out_*) respectively, the corresponding basis element mapping between these types has shape (*o_in_, o_in_* · *o_out_*). Let *f* = min (*o_in_, o_out_*) denote the frequency of the mapping, then for each pair of features we require a radial kernel of size (*d_in_, d_out_, f*). To ensure equivariance, the kernel passes through the corresponding basis element, yielding an intermediate tensor of shape

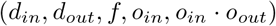

That is, the kernel is obtained from multiplying the radial weights for each pair through the corresponding basis element. The input features are then multiplied through the respective kernel to yield the desired output, and the process is repeated for each pair of input and output types.

As TFNs are used to produce key and value vectors in each attention block, the intermediate kernel mapping between type-ℓ features at each block requires memory proportional to

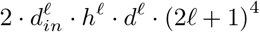

where *h*^ℓ^, *d*^ℓ^ are are the number of heads and head dimension for type ℓ features.

In our implementation, we are able to obtain a factor ≈ *o_in_* · *o_out_* reduction in memory by changing change the order of matrix multiplication in the TFN kernel. Rather than multiply the radial weights through the basis, we first multiplying the features through the basis, and then multiplying the result with the radial weights. The memory required to store the intermediate tensors is reduced to

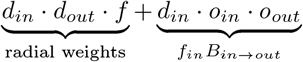

This greatly reduces the memory burden of TFNs and, together with gradient checkpointing, allows us to fit a much deeper and larger model on a single GPU.

### S9 Extended Results

#### S9.1 CASP-FM RMSD

To better understand AttnPacker’s ability to generalize to new folds, we evaluated each method on CASP13 and CASP14 free modelling (FM) targets. These datasets consist of proteins with previously unseen folds and hard analogous fold based models (see Table S11 for a complete list). The results are show in Table S6.

**Table S6:** Average RMSD (Å) on the CASP13-FM and CASP14-FM targets. Results are divided by residue centrality (All, Core, and Surface).

| Method | RMSD (Å) |  |  |  |  |  |
| --- | --- | --- | --- | --- | --- | --- |
|  | CASP13-FM |  |  | CASP14-FM |  |  |
|  | All | Core | Surface | All | Core | Surface |
| SCWRL | 0.741 | 0.441 | 0.919 | 1.008 | 0.681 | 1.206 |
| FASPR | 0.726 | 0.468 | 0.896 | 0.999 | 0.678 | 1.204 |
| RosettaPacker | 0.701 | 0.372 | 0.892 | 0.963 | 0.562 | 1.164 |
| DLPacker | 0.617 | 0.413 | 0.776 | 0.903 | 0.618 | 1.094 |
| TFN-Transformer | 0.591 | 0.431 | 0.705 | 0.850 | 0.629 | 0.987 |
| +Tri | **0.538** | **0.347** | **0.664** | <u>0.784</u> | **0.530** | <u>0.944</u> |
| +Tri+Local | <u>0.542</u> | <u>0.349</u> | <u>0.671</u> | **0.779** | <u>0.531</u> | **0.943** |
| Residue Count | 5465 | 1672 | 2503 | 3934 | 915 | 2012 |

#### S9.2 Non-Native Backbone

**Table S7:** Average RMSD and Dihedral results by method for CASP14-FM non-native backbone targets produced by AlphaFold2. RMSD is shown for all, core, and surface (sfc) residues in angstroms. Overall χ_1–4_ MAE is shown for each methods in units of degrees. Last, overall accuracy for χ_1–4_ angle predictions is shown for all, core, and surface residues.

| | RMSD (Å) | | | MAE° | | | | $\chi$ - Accuracy | | |
| --- | --- | --- | --- | --- | --- | --- | --- | --- | --- | --- |
| | All | Core | Sfc | $\chi_1$ | $\chi_2$ | $\chi_3$ | $\chi_4$ | All | Core | Sfc |
| SCWRL | 1.203 | 0.919 | 1.360 | 41.51 | 34.46 | 48.02 | <u>46.59</u> | 52.4% | 63.4% | 47.3% |
| FASPR | 1.194 | 0.896 | 1.359 | 41.54 | 33.93 | 49.12 | 47.67 | 51.8% | 62.5% | 46.3% |
| Ros. Pack | 1.178 | 0.883 | 1.339 | 40.88 | 34.24 | 47.89 | 48.32 | 53.0% | 63.0% | <u>47.7%</u> |
| DLPacker | 1.138 | 0.857 | 1.299 | 40.51 | 34.74 | 52.46 | 66.49 | 49.3% | 62.2% | 41.9% |
| AlphaFold2 | 1.080 | 0.826 | 1.216 | **37.84** | 30.76 | **42.21** | **44.01** | **55.7%** | **65.8%** | **50.8%** |
| TFN+Tri | 1.102 | 0.874 | 1.226 | 39.75 | **30.20** | <u>47.86</u> | 50.92 | 52.5% | 63.2% | 46.7% |
| TFN+Tri(25) | **1.046** | <u>0.817</u> | **1.170** | <u>38.99</u> | <u>30.35</u> | 48.10 | 48.58 | <u>53.9%</u> | <u>65.4%</u> | 46.2% |
| TFN+Tri(75) | <u>1.059</u> | **0.813** | <u>1.184</u> | 39.59 | 30.46 | 48.65 | 49.12 | 53.8% | 65.9% | 46.2% |

For non-native backbone comparison we trained two separate models TFN+Tri(25) and TFN+Tri(75). These models were trained using the same list of targets as the original model, but with each target having backbone coordinates predicted by AlphaFold2 in place of the native with probability 25% and 75% respectively.

To train these models, we calculated separate rotations and translations mapping the backbone atom coordinates of each residue in the native structure to the corresponding coordinates of the decoy (non-native) structure. We then applied these transformations to align both backbone and side-chain coordinates for each residue in the native structure and treated the result as the ground-truth. After applying this transformation, we were able to use the loss functions described in Section S3.1 without modification.

The average RMSD results shown in Section S9.2 were calculated by aligning the native and decoy residue coordinates as described in the previous paragraph. Substituting non-native backbone coordinates with 25% probability yields the best results in terms of Average RMSD, even better than that of AlphaFold2. Both models, trained partially with non-native backbone data, outperform the baseline TFN+Tri model for side-chain RMSD minimization. In terms of side-chain dihedral prediction accuracy, Alphafold2 achieves the best results, with our methods achieving competitive scores in accuracy for all but surface residues. Similar to the results on native backbones, our models are competitive in terms of dihedral MAE for χ_1_ and χ_2_, but lose their edge for higher order χ values.

#### S9.3 Per Residue RMSD

**Table S8:** Average Per-Residue RMSD for CASP13 Targets by method (column) and residue type (row). Here, TFN+Loc denotes the TFN transformer with local triangle updates. TFN+Tri denotes the TFN Transformer with global triangle updates.

|  | RMSD (Å) |  |  |  |  |  |  | Res.<br>count |
| --- | --- | --- | --- | --- | --- | --- | --- | --- |
|  | SCWRL4 | FASPR | Rosetta<br>Packer | DLPack | TFN | TFN+<br>Loc | TFN+<br>Tri |  |
| ARG | 2.119 | 2.086 | 1.938 | 1.802 | 1.762 | 1.572 | **1.535** | 1040 |
| ASN | 0.918 | 0.889 | 0.873 | 0.761 | 0.751 | 0.706 | **0.671** | 1070 |
| ASP | 0.882 | 0.903 | 0.875 | 0.723 | 0.693 | **0.616** | <u>0.645</u> | 1153 |
| CYS | 0.517 | 0.543 | 0.409 | 0.432 | 0.382 | 0.331 | **0.307** | 240 |
| GLN | 1.456 | 1.450 | 1.303 | 1.217 | 1.163 | 1.086 | **1.077** | 815 |
| GLU | 1.460 | 1.451 | 1.400 | 1.288 | 1.207 | 1.137 | **1.111** | 1192 |
| HIS | 0.989 | 0.865 | 0.855 | 0.702 | 0.686 | 0.679 | **0.605** | 446 |
| ILE | 0.575 | 0.576 | 0.548 | 0.524 | 0.514 | 0.470 | **0.457** | 1238 |
| LEU | 0.607 | 0.595 | 0.571 | 0.521 | 0.513 | 0.463 | **0.456** | 1993 |
| LYS | 1.619 | 1.592 | 1.521 | 1.400 | 1.315 | **1.210** | <u>1.219</u> | 1100 |
| MET | 1.334 | 1.228 | 1.172 | 1.122 | 1.166 | 0.976 | **0.952** | 481 |
| PHE | 0.816 | 0.715 | 0.730 | 0.545 | 0.586 | **0.457** | <u>0.459</u> | 921 |
| PRO | 0.253 | 0.239 | 0.233 | 0.190 | 0.208 | **0.190** | <u>0.201</u> | 943 |
| SER | 0.711 | 0.713 | 0.695 | 0.583 | 0.568 | 0.550 | **0.523** | 1480 |
| THR | 0.508 | 0.497 | 0.477 | 0.440 | 0.409 | **0.387** | <u>0.395</u> | 1317 |
| TRP | 1.312 | 1.128 | 0.981 | 0.830 | 0.995 | 0.700 | **0.695** | 374 |
| TYR | 1.046 | 0.967 | 1.000 | 0.672 | 0.747 | 0.626 | **0.601** | 860 |
| VAL | 0.316 | 0.315 | 0.307 | 0.283 | 0.285 | 0.265 | **0.261** | 1476 |

**Table S9:**
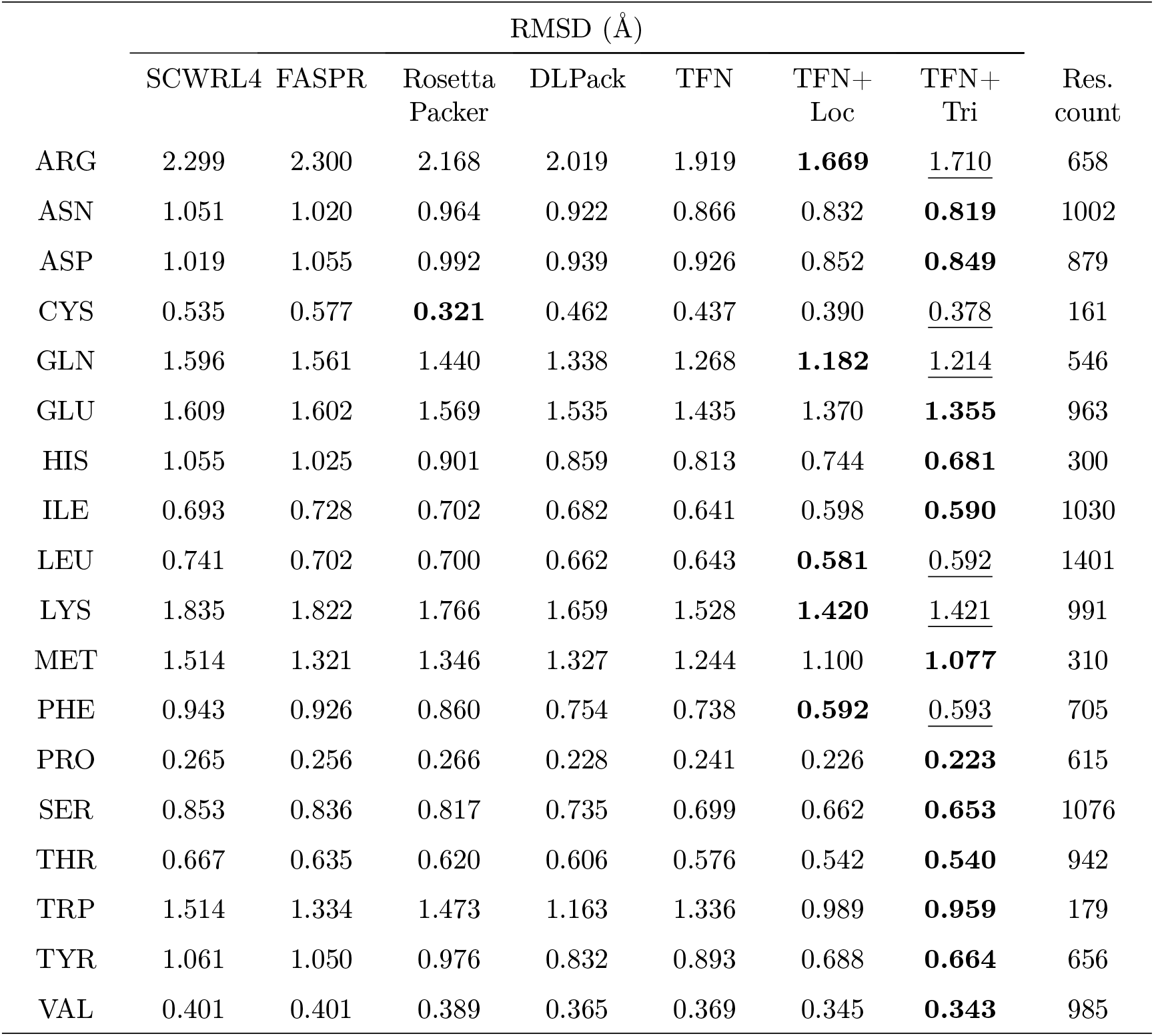
Average Per-Residue RMSD for CASP14 Targets by method (column) and residue type (row). Here, TFN+Loc denotes the TFN transformer with local triangle updates. TFN+Tri denotes the TFN Transformer with global triangle updates.

**Table S10:** Dihedral MAE results on CASP14 targets for charged and polar Amino Acids with high degrees of freedom.

| Method | MAE° |  |  |  |  | MAE° |  |  |  |
| --- | --- | --- | --- | --- | --- | --- | --- | --- | --- |
| | $\chi_1$ | $\chi_2$ | $\chi_3$ | $\chi_4$ | acc. | $\chi_1$ | $\chi_2$ | $\chi_3$ | acc. |
|  | ARG |  |  |  |  | GLN |  |  |  |
| SCWRL | 38.97 | 33.5 | 67.92 | 57.25 | <u>49.0%</u> | 36.81 | 48.3 | 39.1 | 47.8% |
| FASPR | 38.85 | 33.87 | 68.97 | 65.05 | 48.5% | 34.93 | 46.16 | 37.45 | 50.8% |
| RosettaPacker | 37.42 | 35.93 | **61.1** | <u>56.99</u> | **50.5%** | 32.38 | 40.96 | **35.78** | <u>53.0%</u> |
| DLPacker | **31.17** | 35.04 | 72.25 | 68.41 | 45.3% | **29.69** | 37.11 | 38.41 | 52.3% |
| TFN-Transformer | 35.71 | 32.20 | 72.67 | 56.92 | 42.0% | 31.75 | 39.03 | 36.94 | 51.1% |
| +Tri | 33.78 | <u>32.58</u> | <u>61.4</u> | **56.03** | 47.8% | 30.52 | **35.66** | 36.48 | **54.0%** |
| +Tri+Local | <u>33.15</u> | **31.35** | 62.57 | 57.23 | 44.8% | <u>30.47</u> | <u>36.32</u> | <u>36.35</u> | 52.1% |
|  | LYS |  |  |  |  | GLU |  |  |  |
| SCWRL | 37.85 | 32.86 | **43.65** | 47.74 | 55.8% | 42.05 | 42.02 | 39.35 | 46.8% |
| FASPR | 37.55 | 31.76 | <u>44.16</u> | **46.39** | **56.0%** | 39.47 | 40.45 | 41.17 | <u>48.0%</u> |
| RosettaPacker | 37.38 | 33.62 | 44.45 | 47.83 | <u>55.1%</u> | 40.24 | 40.45 | 39.34 | <u>48.0%</u> |
| DLPacker | 32.48 | 33.94 | 50.34 | 68.32 | 45.9% | 39.37 | 42.24 | 40.43 | 45.1% |
| TFN-Transformer | 29.59 | 33.61 | 44.60 | 48.83 | 50.8% | 36.24 | 39.61 | 36.97 | 46.8% |
| +Tri | **28.04** | **29.86** | 44.9 | 47.89 | 54.2% | **34.37** | **37.25** | **35.86** | **50.5%** |
| +Tri+Local | <u>29.12</u> | <u>30.91</u> | 44.34 | <u>46.79</u> | 53.0% | <u>36.71</u> | <u>37.39</u> | <u>36.17</u> | 47.1% |

### S10 PDB Lists

**Table S11:** List of targets in each test data set.

| Data Set | Targets |
| --- | --- |
| CASP13 | T0949, T0950, T0951, T0953s1, T0953s2, T0954, T0955, T0957s1, T0957s2, T0958, T0959, T0960, T0961, T0962, T0963, T0964, T0965, T0966, T0967, T0968s1, T0968s2, T0969, T0970, T0971, T0973, T0974s1, T0974s2, T0975, T0976, T0977, T0978, T0979, T0980s1, T0980s2, T0981, T0982, T0983, T0984, T0985, T0986s1, T0986s2, T0987, T0988, T0989, T0990, T0991, T0992, T0993s1, T0993s2, T0994, T0995, T0996, T0997, T0998, T1000, T1001, T1002, T1003, T1004, T1005, T1006, T1008, T1009, T1010, T1011, T1013, T1014, T1015s1, T1015s2, T1016, T1016_A, T1017s1, T1017s2, T1018, T1019s1, T1019s2, T1020, T1021s1, T1021s2, T1021s3, T1022s1, T1022s2 |
| CASP14 | T1024, T1025, T1026, T1027, T1028, T1030, T1031, T1032, T1033, T1034, T1035, T1036s1, T1037, T1038, T1039, T1040, T1042, T1043, T1045s1, T1045s2, T1046s1, T1046s2, T1047s1, T1047s2, T1048, T1049, T1052, T1053, T1054, T1055, T1056, T1057, T1058, T1060s2, T1060s3, T1062, T1064, T1065s1, T1065s2, T1067, T1068, T1070, T1072s1, T1073, T1074, T1078, T1079, T1080, T1082, T1083, T1084, T1087, T1088, T1089, T1090, T1091, T1092, T1093, T1094, T1095, T1096, T1098, T1099, T1100 |
| CASP13-FM | T0950, T0953s1, T0953s2, T0957s1, T0957s2, T0960, T0963, T0968s1, T0968s2, T0969, T0975, T0980s1, T0981, T0986s2, T0987, T0989, T0990, T0991, T0998, T1000, T1001, T1010, T1015s1, T1017s2, T1021s3, T1022s1 |
| CASP14-FM | T1027, T1031, T1033, T1037, T1038, T1039, T1040, T1042, T1043, T1047s1, T1049, T1064, T1070, T1074, T1090, T1093, T1094, T1096 |

